# A unifying perspective of the ultrafast photo-dynamics of Orange Carotenoid Protein from *Synechocystis*: peril of high-power excitation, existence of different S* states and influence of tagging

**DOI:** 10.1101/2021.12.26.474187

**Authors:** Stanisław Niziński, Adjéle Wilson, Lucas M. Uriarte, Cyril Ruckebusch, Elena A. Andreeva, Ilme Schlichting, Jacques-Philippe Colletier, Diana Kirilovsky, Gotard Burdzinski, Michel Sliwa

## Abstract

A substantial number of Orange Carotenoid Protein (OCP) studies have aimed to describe the evolution of singlet excited states leading to the formation of photo-activated form, OCP^R^. The most recent one suggests that three picosecond-lived excited states are formed after the sub-100 fs decay of the initial S_2_ state. The S* state which has the longest reported lifetime of a few to tens of picoseconds is considered to be the precursor of the first red photoproduct P_1_. Here, we report the ultrafast photo-dynamics of the OCP from *Synechocystis* PCC 6803, carried out using Visible-NIR femtosecond time-resolved absorption spectroscopy as a function of the excitation pulse power and wavelength. We found that a carotenoid radical cation can form even at relatively low excitation power, obscuring the determination of photo-activation yields for P_1_. Moreover, the comparison of green (540 nm) and blue (470 nm) excitations revealed the existence of an hitherto uncharacterized excited state, denoted as S^∼^, living a few tens of picoseconds and formed only upon 470 nm excitation. Since neither the P_1_ quantum yield nor the photo-activation speed over hundreds of seconds vary under green and blue continuous irradiation, this S^∼^ species is unlikely to be involved in the photo-activation mechanism leading to OCP^R^. We also addressed the effect of His-tagging at the N- or C-termini on excited state photo-physical properties. Differences in spectral signatures and lifetimes of the different excited states were observed, at variance with the usual assumption that His-tagging hardly influences protein dynamics and function. Altogether our results advocate for careful consideration of the excitation power and His-tag position when comparing the photo-activation of different OCP variants, and beg to revisit the notion that S* is the precursor of photoactivated OCP^R^.

**TOC:** 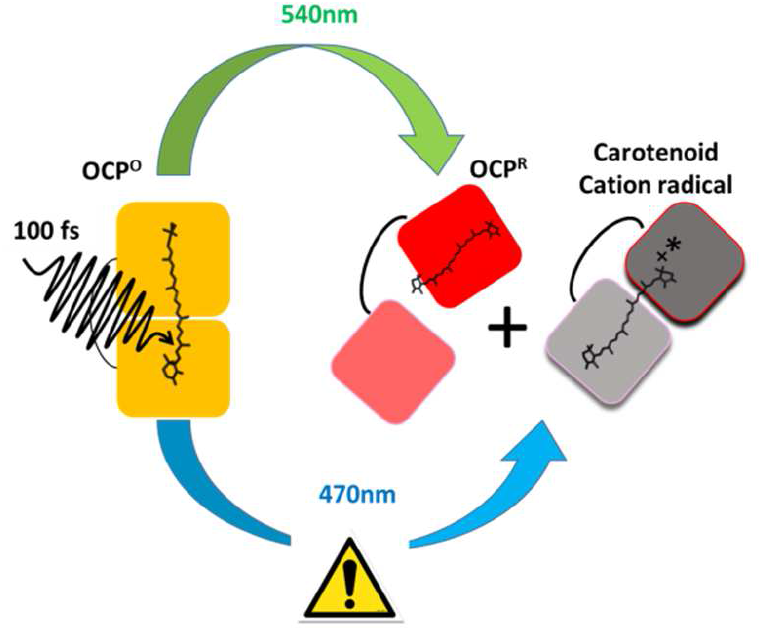

## Introduction

The Orange carotenoid protein (OCP) is a 35 kDa water-soluble photo-active protein capable of quenching the excess of light energy harvested by cyanobacteria.^1-4^ In order to perform its energy-quenching function, dark adapted OCP (abbreviated as OCP^O^ due to its orange color) must be photo-activated by strong blue-green light illumination, yielding the OCP^R^ species capable of quenching excited phycobilisomes.^5^ The protein is structured as a two-domain protein, where the N-terminal (NTD) domain is the effector, and the C-terminal (CTD) domain is the regulator.^6-7^ The functionalizing ketocarotenoid chromophore is embedded at the interface between the NTD and CTD.^8^ The main function of OCP is to quench the fluorescence of the cyanobacterial light harvesting antennas, a.k.a. phycobilisomes. Only the OCP^R^ state can interact with the latter and perform the quenching function.

The photo-activation of OCP^O^ starts with the evolution of the chromophore excited-state levels resulting in the formation of P_1_. This state is the first red photoproduct with broken H-bonds between the carotenoid and the protein.^9^ The 12 Å migration of the ketocarotenoid in the NTD^10^ and structural changes in the protein, respectively occurring in the microsecond and millisecond time scale, ultimately^9, 11-12^ lead to separation of the two domains (Scheme 1a), yielding OCP^R^. The photo-conversion quantum yield of OCP^R^ is very low (0.2% or less)^5, 9, 11-12^, reflecting the functional requirement that OCP remains inactive in low irradiance conditions, i.e. when maximum energy transfer to photochemical centers is needed. The low OCP photo-conversion quantum yield is determined during excited-state deactivation, with about 99% of the ketocarotenoid relaxing back to the initial S_0_ state within tens of picoseconds.^5, 9, 12-13^ The ultrafast photo-dynamics of different OCPs and carotenoids (hydroxyechinenone, echinenone, canthaxanthin, zeaxanthin) have been studied by femtosecond transient absorption spectroscopy using various excitation wavelengths in the visible range.^5, 9, 14-18^ The most recent studies consider that upon relaxation of the initial S_2_ state, three picosecond-lived excited states are formed: a sub-picosecond-lived Intramolecular Charge Transfer (ICT) state, a picosecond-lived mixed S_1_/ICT state called usually S_1_, and an excited state S* characterized by a lifetime in the range of few picoseconds^13^ to few tens of picoseconds^9^, depending on the source. There is a longstanding debate regarding the ground vs. excited state nature of the carotenoid S* state, but no agreement has been reached thus far.^19-22^ The carotenoid S* was first postulated to be a hot ground state,^19^ then redefined as an electronically excited state,^20^ but some publications questioned this hypothesis and pointed out different properties featuring a vibrationally hot electronic ground state.^21-24^ Regardless, the ICT, S_1_ and S* populations are all known to decay within few tens of picoseconds, while the first red photoproduct P_1_, appears with a yield of about 1.5%.^9^ Despite this already vast knowledge, the low magnitude of the P_1_ signal implies that a high error intrinsically exists on the determined quantum yield. Furthermore, it remains unclear which of the excited states are the precursor of P_1_. Recent studies have considered S*, on the basis that S* would have a distorted geometry which favors the breaking of hydrogen bond and formation of P_1_.^9^ However, the issues concerning the genuine nature of S* render difficult any firm conclusion.

**Scheme 1:**
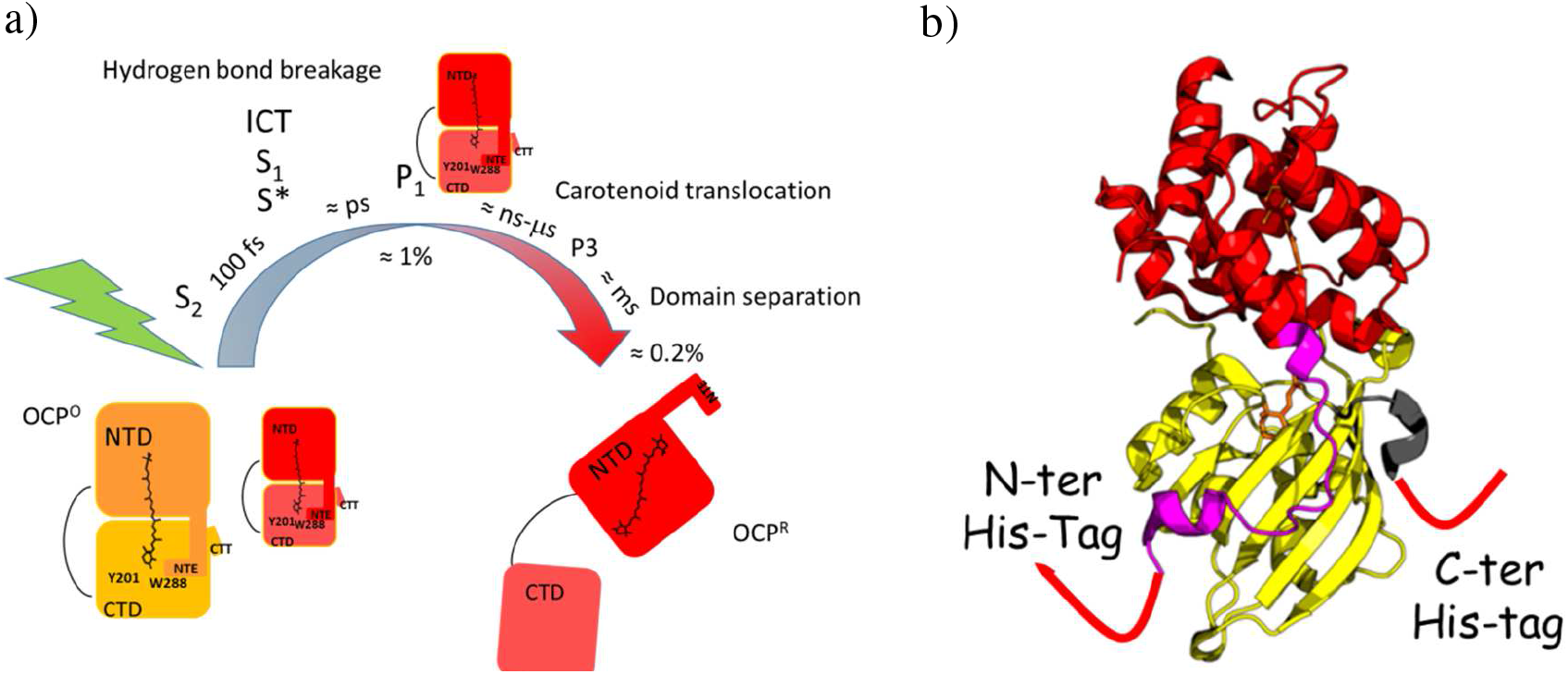
a) General photo-dynamical scheme of OCP photo-activation mechanism. In the dark, two subpopulations of the closed OCP^O^ are present, a “normal” one and a redshifted one (the size represents their contribution, Figure S2 in Supporting Information). b) The OCP structure (reproduced from PDB ID 3MG1) comprises a fully α-helical NTD, featuring a fold that is unique to cyanobacteria, whereas the CTD has a mixed α/β architecture and belongs to the NTF-2 family.^8^ His-tags can be attached to the N-terminal extension (NTE) or to the C-terminal tail (CTT).

Kennis and coworkers were the first to report that the formation yield and lifetime of the S* state depend on the excitation peak power.^9^ Such results could be explained by the existence of additional pathways and extra species, possessing a similar spectra but different lifetimes, and generated by multiphotonic processes caused by femtosecond pulse excitation – as reported for other photo-active proteins.^25-26^ Multiphotonic processes are usually accompanied by off-pathway species, such as radical cations and solvated electrons, which are avoided when the excitation pulse is stretched or the excitation energy reduced.^25-26^ The Polivka group showed an increase of the S* population and the existence of a long lived radical signal peaking at 900 nm, when probing the effect of UV excitation on canthaxantin-functionalized OCP.^27^ A functional role of an oxocarbenium cation was even suggested in the photo-activation of OCP^O^.^13^ Surprisingly, however, despite the fact that the low energy keto-carotenoid absorption band is associated with the S_0_→S_2_ transition, no studies have thus far investigated the power dependence of excitation using femtosecond pulses. Clearly, knowledge of the photoexcitation power dependence and characterization of radical cation is crucial for the interpretation of OCP ultrafast dynamics. Also, two excitation wavelengths have been used depending on studies, viz. 470 nm and 540 nm,^14, 16, 28^ due to existence of at least two ground state populations in the dark state (referred to as “normal” and “redshifted”^14, 29^). The 540 nm excitation presumably selects only the redshifted subpopulation, while 470 nm excites both of them. Last, studies have been performed on the C-tagged or N-tagged OCPs,^5, 9, 14-18^ yet the (admittedly unlikely) hypothesis that tagging could influence ultrafast photo-dynamics was never investigated, despite both tag being located on helices attached to the sensory domain, *i*.*e*. the CTD (Scheme 1). It could thus be that tagging affects the equilibrium between the normal and red-shifted dark-adapted OCP^O^ (Scheme 1) and/or their excited-state dynamics, which could in turn influence the photo-conversion yield.

Here we report a detailed photo-physical study on OCP from *Synechocystis* PCC 6803 complexed with the ketocarotenoid echinenone (ECN). The use of visible-NIR femtosecond transient absorption spectroscopy allowed to identify all species involved in the OCP excited state deactivation. With view to afford comparison with all previously published transient spectroscopy studies,^5, 9, 14-18^ we investigated light-induced excited state dynamics upon excitation by either 470 nm or 540 nm light. Stationary irradiation experiments were also performed at the two excitation wavelengths. Both on the C-tagged or N-tagged variant of OCP were studied, to test the effect of the tag and its position on ultrafast photo-dynamics. On each construct and at the two wavelengths, we furthermore probed the effect of the excitation power on excited state dynamics. We show how the picosecond dynamics that determine formation of P_1_, the first crucial intermediate controlling the formation quantum yield of OCP^R^, are influenced by the excitation energy and wavelength. The combination of transient absorption and stationary irradiation experiments upon 470 nm and 540 nm excitation allows us to identify which of the picosecond-lived states correlates with the final photo-activation quantum yield, and is thus the most likely candidate to be the precursor of P_1_. We show that the use of high excitation power leads to the formation of a carotenoid radical cation, whose presence may result in an erroneous extraction of P_1_ yield from OCP bleach kinetics. Finally, we demonstrate that upon 470 nm excitation, the photo-dynamics of OCP feature a yet unidentified S^∼^ state whose existence may compromise estimation of the S* lifetime and P_1_ yield. Taking into account all the above-mentioned aspects of OCP photoexcitation will prove crucial, not only to understand the basis for the biological function of OCP, but also to design of OCPs with a higher photo-activation quantum yield.

## Materials and Methods

### Protein expression and purification

The plasmid pCDF-NtagOCPSyn used for expression of OCP from *Synechocystis* PCC 6803 carries a sequence coding for a His-tag in the N terminus, and was characterized in Bourcier de Carbon et al.^30^ The expression of the *ocp* genes in *E. coli* cells containing the genes for synthesis of echinenone and the isolation of ECN-OCP was also described.^30^ The expression of the *ocp* genes in *Synechocystis* was reported by Gwizdala et al.^31^ N-tagged OCP was expressed in *E. coli*., while C-tagged OCP was expressed in the ΔcrtR *Synechocystis* mutant lacking zeaxanthin and hydroxyechinenone. The protein concentration used in the experiments described in this work was close to 1 mg/ml (calculated using absorbance at 496nm) and 40 mM Tris-HCl 25 mM NaCl pH 8.0 buffer was used.

### Steady state UV-vis absorption spectra and photoconversion under steady state irradiation

Steady state UV-vis absorption spectra were recorded with a Jasco V-550 spectrophotometer, using 2 nm spectral bandwidth. The cuvette optical path was 10 mm for stationary LED irradiation spectroscopy experiments. The photo-conversion kinetics set up is described elsewhere.^32^ Here the photo-activation was induced by green LED irradiation (λ_max_=528 nm, FWHM=28 nm, standard 3W emitter), and blue LED irradiation (λ_max_=470 nm, FWHM=18 nm, standard 3W emitter). The probing and irradiation beams were at 90°. The optical irradiation path of the solution was 4 mm; the probing path was 10 mm. The absorbance at 470 nm and 528 nm were the same for C-tagged and N-tagged OCP (0.53 and 0.28 for 470 nm and 528 nm, respectively). The irradiation power was 3.4 mW/cm^2^ for the blue LED and 3.0 mW/cm^2^ for the green LED, therefore the photon flux is the same in both cases.

### Femtosecond Vis-NIR transient absorption

Femtosecond Vis-NIR transient absorption spectra were collected using a commercially available system (Ultrafast Systems, Helios) described previously^33^ that consists of a short-pulse titanium-sapphire oscillator (Mai-Tai, Spectra Physics, 70 fs) followed by a high-energy titanium-sapphire regenerative amplifier (Spitfire Ace, Spectra Physics, 100 fs, 1 kHz). The 800 nm beam was split into two beams to generate: (1) the pump (λ_exc_ = 540 nm or 470 nm) in the optical parametric amplifier (Topas Prime with a NirUVVis frequency mixer) and (2) probe pulses - white light continuum in the Vis-NIR range generated by focusing the fundamental beam into a sapphire (430–780 nm), or YAG (820–1390 nm) crystal. The remaining 800 nm probe pulse photons were filtered. The instrument response function (IRF) was determined by fitting the kinetics of the coherent artefact signal from solvent and was estimated to be ≈110 fs (FWHM). The experiments were performed with different pulse energies ranging from 0.2 μJ (3.3 × 10^14^ photons per cm^2^ at FWHM, Figure S5) up to 1.6 μJ upon 470 nm excitation and from 0.4 μJ (6.6 × 10^14^ photons per cm^2^ at FWHM, Figure S6) up to 3.2 μJ upon 540 nm excitation using variable neutral density filters. The pump diameter (FWHM) at the sample was ≈250 µm. Number of photon and fluence per pulse are given in the supporting information (Fig. S5 and Fig. S6). In all transient absorption experiments, the absorbance was close to 0.7 at the excitation wavelength in a 2 mm optical path. The sample solution was stirred to keep fresh OCP solution in the probed volume. The transient spectra were registered with 1 nm per pixel and by averaging 500 supercontinuum spectra with and without excitation, respectively. The entire set of pump-probe time delay points was repeated four times to ensure data reproducibility, then the data were inspected and averaged. Moreover, to ensure that all datasets are comparable to each other, they were measured in one experimental session, in identical conditions except varied parameters (like pump energy, explicitly given). The pump beam was depolarized to avoid anisotropy effects. The sample temperature was set to 22 °C. The stability of the sample was checked by comparing the stationary UV-vis absorption spectra measured before and after the experiments.

### Data analysis

The transient absorption data were corrected for the chirp of white light continuum based on the given amount of sapphire, water and BK7 glass, which the probe pulse had to pass through. Afterwards, visible and NIR data were merged using a custom procedure.^33^ Low intensity of the probing pulse and presence of relatively strong residual 800 nm beam (even after filtering) resulted in minor artifacts visible in recorded data around 800 nm. For all datasets, the difference absorbance value obtained at the bleaching minimum (in both spectral and temporal dimension) was normalized to -1. Transient spectra were projected onto a 5 nm-spaced grid to get kinetic traces. The comparison of pre-exponential factors at 490 nm (bleaching band) allowed us to estimate the formation quantum yield of the various intermediates. Data processing to determine time constants and Decay Associated Difference Spectra (DADS or DAS) was based on our custom fitting procedure. It consists of two steps. Firstly it fits globally representative wavelengths (480, 490, 500, 570, 590, 610, 655, 740, 960, 1100 nm) with convolution (IRF 110fs FWHM) and weights (to increase contribution of the long delays, which are our main interest, to the χ^2^ error term). Secondly, the extracted time constants are fixed and used to fit all kinetics separately, and finally DAS are built. Bootstrapping analysis was also performed to estimated errors (see SI for more details). We also tested our results against other procedures, like the one implemented in the Glotaran package.^34^ For the global analysis a sum of 4 or 5 exponentials convoluted with a Gaussian-shaped pulse of 110 fs (FWHM) and an offset which represents long-lived photo-product(s) (> 10 ns, P_1_ + radical) was used, for 540 nm and 470 nm excitation respectively.

## Results

### Steady state properties

In the OCP^O^ (dark-adapted) state, UV-Visible absorption spectra of N-tagged and C-tagged OCP show almost no difference, both display a broad absorption band, characteristic of the S_0_-S_2_ transition, with two maxima at 472 and 496 nm and a tail until 650 nm (Figure 1a, Figure S1 Supporting Information).^29, 35^ The “redshifted” population was estimated by Gaussian decomposition to be around 20% for both N-tagged and C-tagged OCPs (Figure S2), in line with previous reports.^14^ To determine if a difference in the ability to photoactivate exists between the two dark-adapted subpopulations, N-tagged and C-tagged OCP solutions (same absorbance) were irradiated with blue (470 nm LED) and green (528 nm LED) light, thereby selecting either the “normal” or the “redshifted” subpopulations of OCP^O^. The initial slope of the evolution of the absorbance at 550 nm (Figure 1b), which probes the formation of OCP^R^, shows that both subpopulations of N-tagged OCPs photo-activate with the same efficiency when using different irradiation wavelengths. There is a slight difference in the photo-stationary state (achieved approximately after 600 s of LED irradiation), suggesting that 528 nm irradiation ultimately generates quantitatively more OCP^R^. This result can be explained by a deviation from dark-adapted equilibrium between subpopulations in OCP^O^ state, caused by the extended period of irradiation (e. g. 528 nm selects one subpopulation, but photoactivated OCPs may repopulate both subpopulations after back-conversion). Unexpectedly, the comparison of the evolution of the absorbance traces at 550 nm for the different tags (Figure 1c) reveals a much faster photo-activation of N-tagged OCP than C-tagged OCP (the initial slope is 3.5 times larger for the N-tagged OCP). This difference in OCP^R^ yield could stems either from different excited states dynamics, leading to different P_1_ formation quantum yield, or from changes in the yield of subsequent ground state species forming in the ns-ms time scale (Scheme 1, carotenoid translocation and domain separation dynamics).^9, 11-12^ To split a difference between the two hypotheses, a more detailed analysis of the early stages of the photo-induced processes is needed.

**Figure 1.**
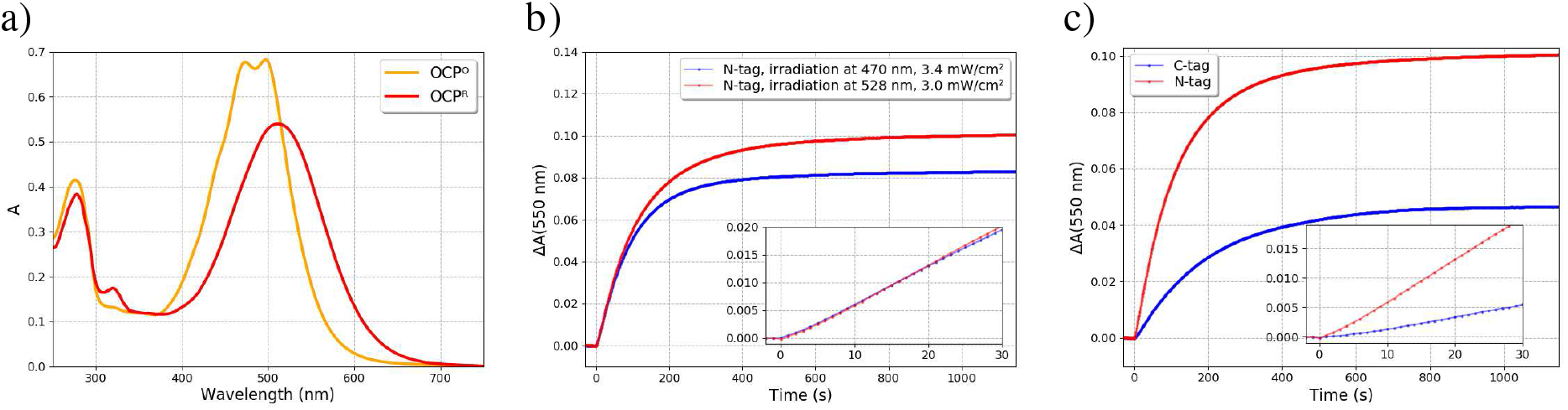
a) UV-vis stationary spectra of OCP^O^ (recorded in the dark) and OCP^R^ (under 452 nm irradiation, 3.2 mW/cm^2^), 1 cm path length, 11°C. b) Evolution of ΔA at 550 nm (22 °C) for N-tagged OCP (N-tag) upon 528 nm (FWHM=28 nm, 3.0 mW/cm^2^) and 470 nm (FWHM=18nm, 3.4 mW/cm^2^) LED irradiation and c) comparison of C-tagged (C-tag) and N-tagged (N-tag) OCP upon 528 nm LED irradiation. Initial slopes are shown in the insets.

### Femtosecond transient absorption spectroscopy of N-tagged OCP

The ultrafast photo-dynamics of N-tagged OCP was studied upon 470 and 540 nm excitation wavelengths. The excitation energy pulse was initially set to 0.4 µJ at 470 nm and to 0.8 µJ at 540 nm, which ensures being in the linear photoexcitation range (Figure 4) while getting enough S/N to characterize P_1_ and keeping the data free of additional signals arising when higher energy is used (detailed discussion in next section). Figure 2 shows femtosecond transient absorption data recorded with 470 nm and 540 nm excitation, respectively. Just after the 540 nm excitation, at 0.05 ps, a primary positive absorption band peaking at 1050 nm is observed. This band is assigned to the S_2_ excited state population (S_2_→S_n_ transition).^17^ The band is broader upon 470 nm excitation and its maximum is shifted to 1100 nm, indicating presence of an excess of vibrational energy and pointing to excitation of a larger range of OCP^O^ subpopulations. Concomitant with formation of the S_2_ state within 50 fs, a ground state bleaching (GSB; indicative of OCP^O^ depopulation) signal is observed in the visible region (see Figure S3 which is zooming into the visible region), with a maximum at about 500 nm, as well as appearance of two main excited-state absorption (ESA) bands, at about 663 nm and 740 nm (Figure 2, Figure S3). These two ESA can be assigned to the S_1_ and ICT states.^14-15^ In addition to the ICT and S_1_ states, the shoulder around 570 nm can be assigned to the S* state in agreement with the recent literature.^9, 13^ After 50 femtoseconds, the first signal evolution is the decay of the S_2_ state within a few hundreds of femtoseconds. During this time, the ESA bands attributed to S_1_ and S* states continue to grow slightly until 0.2 ps. The amplitude of the bands around 570-585 nm (S* contribution) and 663 nm (S_1_ contribution) are higher for the 470 nm excitation. Conversely, and in agreement with previous results,^14-15, 28^ the ICT character is more pronounced when 540 nm excitation is used, resulting in higher amplitude signals around 740 nm and a long tail up to 1100 nm. This increase in amplitude of the ICT character is at the expense of the amplitude of the bands at 663 nm and 570 nm. After 10 picoseconds, the S_1_ and ICT states decay concomitant to recovery of 97% of the GSB band (Figure 2). At 23 ps time delay, the remaining signal is characterized by the broad positive absorption band usually assigned to the S* state, with a maximum at 570 nm and a shoulder around 655 nm (Figure 3).^9^ The comparison of the transient spectra at 23 ps recorded after 470 nm and 540 nm excitations, respectively (Figure 3a and 3b, purple spectrum), reveals major differences, the most notable one being that the GSB band is two times smaller for 540 nm excitation (about 0.008 vs 0.016 for 540 and 470 nm excitations; values estimated from GSB signal at 490 nm). Moreover, while the transient absorption band observed upon 470 nm excitation peaks at 570 nm, that observed upon 540 nm excitation has a much broader shape, with the maximum shifted to ≈585 nm and a much higher amplitude of the shoulder around 655 nm. Thus, both excitations lead to different S* type species. S* state decay within 50 ps, however upon 470 nm excitation there is an extra time evolution of the 570 nm band within 500 ps (Figure 3a). This is assigned to the existence of an additional species S^∼^, forming only upon 470 nm excitation and characterized by a positive absorption band with a maximum located at 570 nm and a lifetime of c.a. 80 ps. Finally, irrespectively of the excitation wavelength, the transient absorption spectrum at 1 ns is characterized by a GSB of OCP^O^ of similar intensity yet a broad positive band also remains, with a maximum at ≈565 nm, which is broader and two times higher in amplitude when a 470 nm pump is used. This band is assigned to P_1_, however only the GSB band, corresponding to OCP molecules that have not relaxed to the initial OCP^O^ state, can afford estimation of the P_1_ yield. Thus using the GSB band of OCP^O^ at 490 nm and taking into account positive absorbance contribution from P_1_ at 490 nm (See Supporting information and Figure S4), P_1_ formation quantum yields is estimated to be ≈ 0.5% for both excitations. These results are also in accordance with the photo-activation efficiency being independent of the irradiation wavelength (470 or 528 nm, Figure 1a).

**Figure 2.**
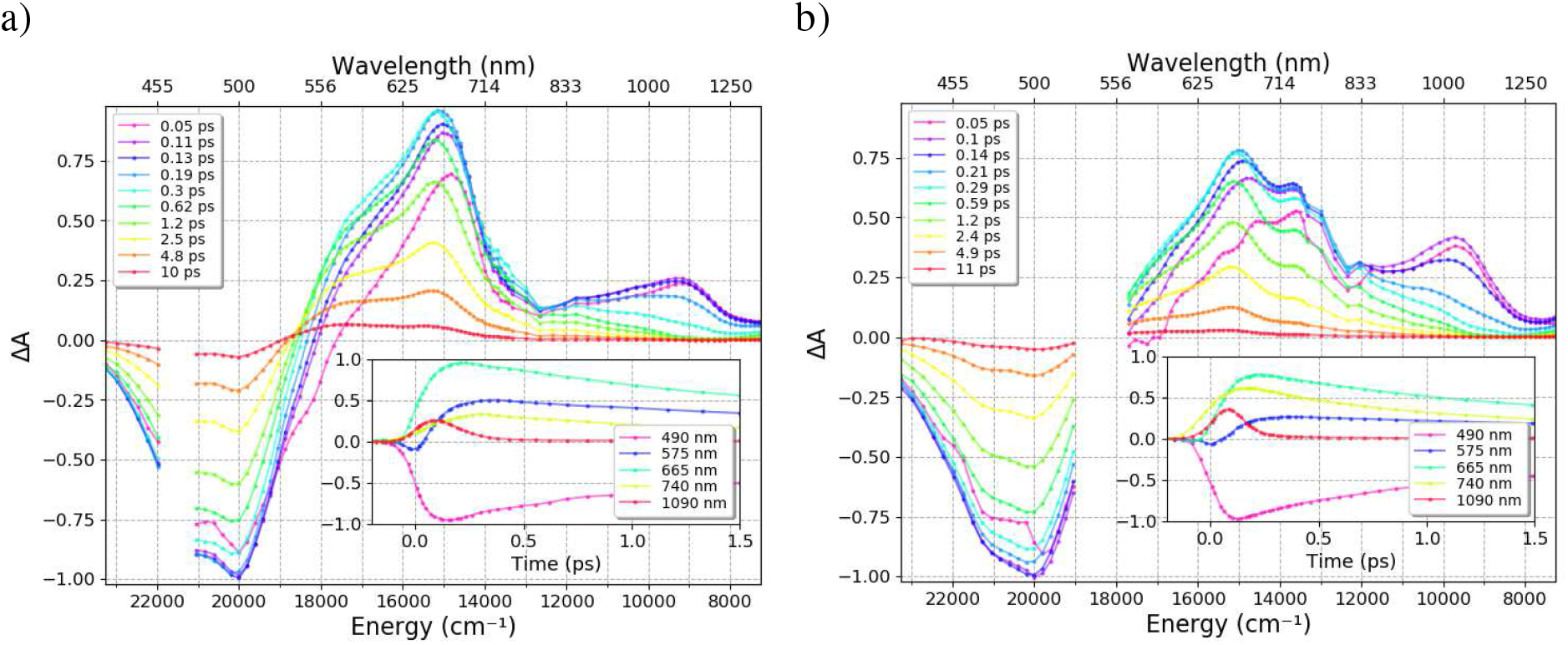
Transient absorption spectra between 0.05 and 10 ps of N-tagged OCP excited at a) 470 nm (0.4 µJ), and b) 540 nm (0.8 µJ). All datasets were normalized to -1 at the bleaching extremum (in both spectral and temporal dimensions). To obtain the original signal, multiply plotted values by a) 0.077 and b) 0.088. Insets show evolution of the signal in the 1.5 ps time window.

**Figure 3.**
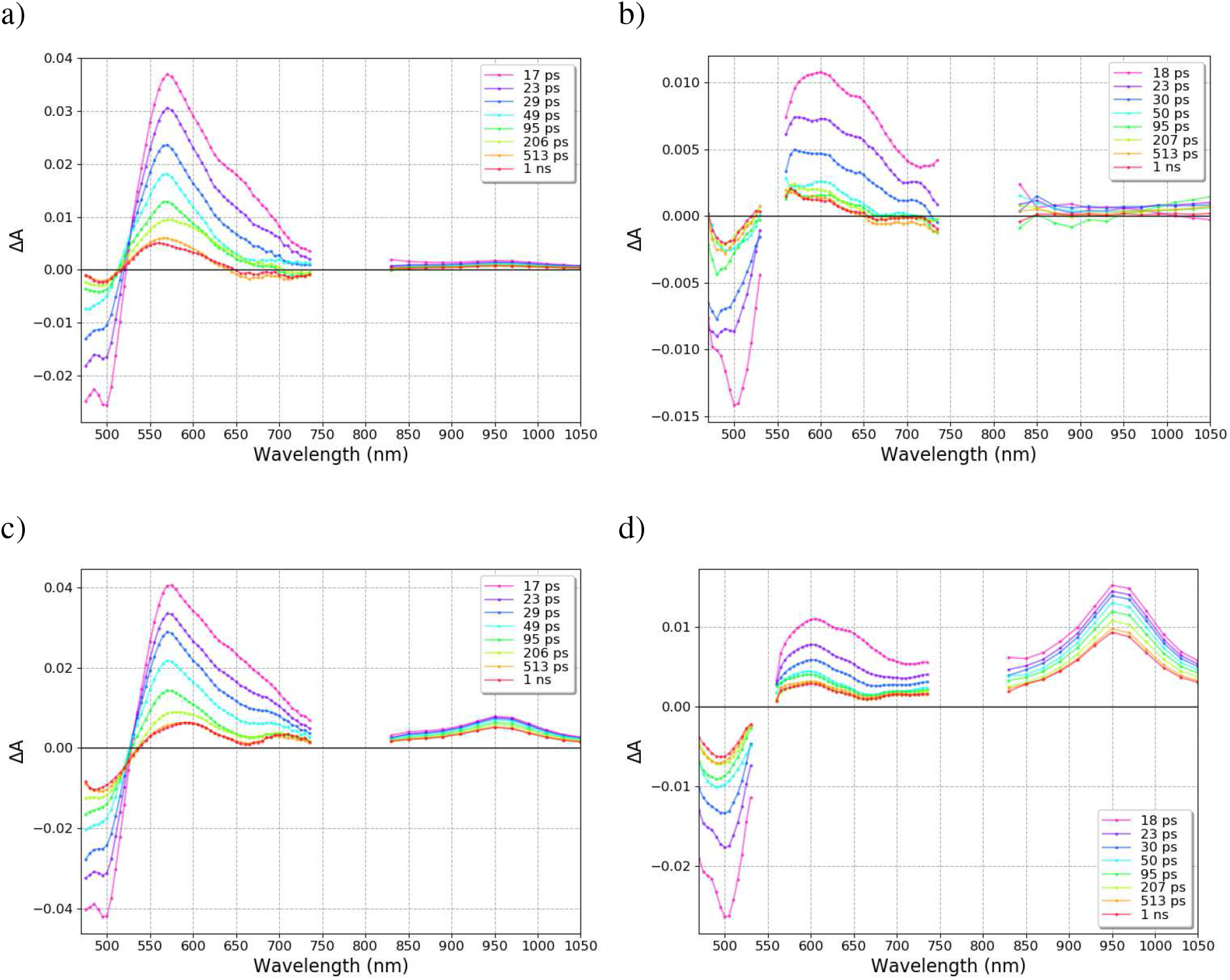
Transient absorption spectra between 17 ps and 1 ns for N-tagged OCP excited at a) 470 nm (0.4 µJ), b) 540 nm (0.8 µJ), c) 470 nm (1.6 µJ), and d) 540 nm (3.2 µJ) All datasets were normalized to -1 at the bleaching extremum (in both spectral and temporal dimensions, multiply by a) 0.077, b) 0.088, c) 0.163 and d) 0.234 to get original signal).

**Figure 4.**
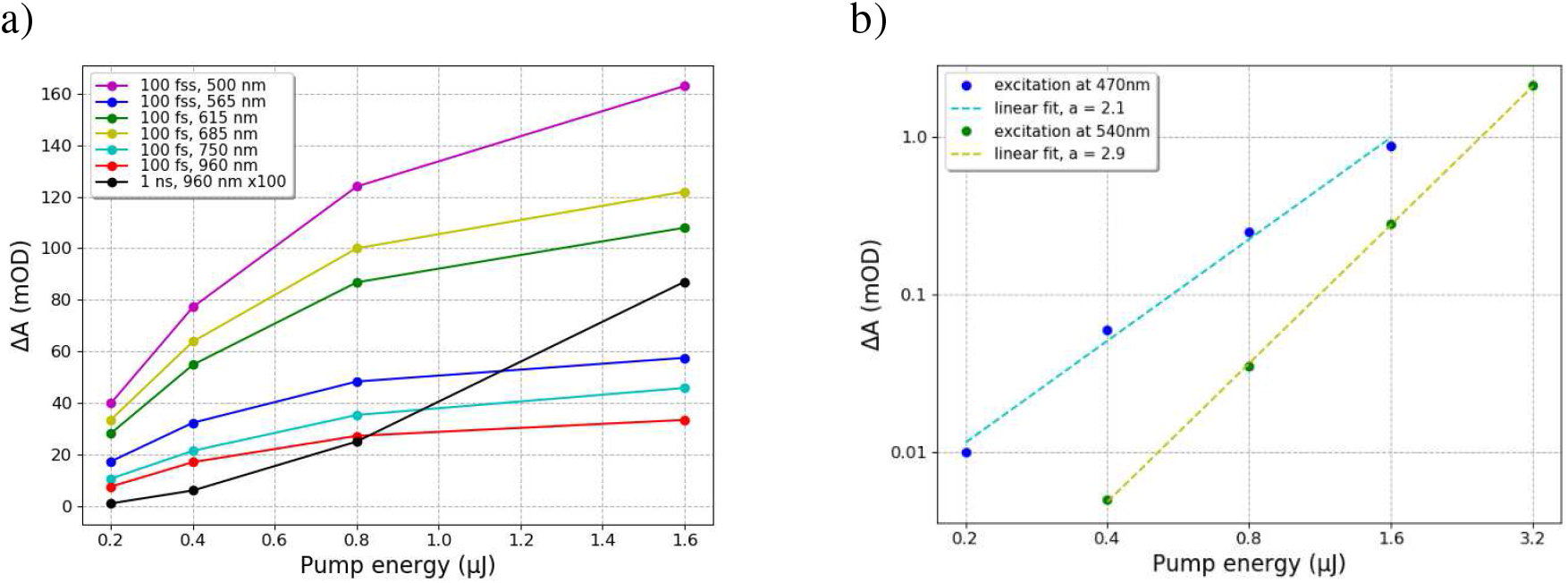
a) Difference absorbance value at different wavelengths versus pump energy for 470 nm excitation (the value at 500 nm was multiplied by -1), b) log-log plots of cation radical absorbance (960 nm) at 1 ns time delay versus excitation energy upon 470 nm and 540 nm excitation.

### Effect of high energy pump pulse excitation

Power dependence measurements were done in the energy range of 0.2-1.6 µJ for 470 nm and 0.4-3.2 µJ for 540 nm excitation. Transient absorption spectra for 1.6 µJ (470 nm) and 3.2 µJ (540 nm) are shown for short time delays in the supporting information (Figure S7) and presented in Figures 3c and 3d for time delays between 17 ps and 1 ns. It should be noted that higher energies were also explored, however fast sample degradation was observed as well as an increase in the scattering of the excitation beam. For the highest energy pulse, the number of photons absorbed per chromophore in the centre of the beam was close to 0.8 (Figures S5 and S6). Figure 3 shows that at 1 ns time delay (Figure 3a and 3b) only P_1_ is present for low excitation energies (positive band with a maximum at 565 nm), while additional positive absorption bands at 700 nm and 960 nm (Figures 3c and 3d), characteristic of radical cation species of carotenoid^36-37^, are observed for high excitation energies. A minor contribution is also present at 600 nm, overlapping with the P_1_ absorbance band (compare Figure 3a with Figure 3c at 1 ns time delay). The presence of a radical cation is characterized by an increase in the depopulation (GSB) band maximum value versus pulse excitation energy (band at 500 nm), while the maximum value of absorbance for S*, S_1_, ICT states (565nm / 685nm / 750 nm) is reaching a plateau for excitation energies exceeding 0.8 µJ for 470 nm and 1.6 µJ for 540 nm (Figure 4). The radical cation signal also affects the P_1_ positive absorption band, although to a lesser extent than the GSB band which peaks to 1% depopulation of OCP^O^ when excitation energy reaches 1.6 µJ upon 470 nm excitation (Figure 3c). The thresholds for the observation of the radical cation are 0.4 and 0.8 µJ at 470 and 540 nm, respectively, with the signal at 960 nm being negligible at these excitation energies (Figure 4). These were therefore selected to characterize the photodynamics of OCP with the best possible S/N ratio, while avoiding contribution of biological irrelevant species. Figures 4b show a log-log plot of absorbance at 960 nm and 1 ns time delay versus excitation energy, underlining the multiphotonic nature of the process leading to the formation of the radical cation. Indeed two photons are involved in the formation of the radical for 470 nm excitation (a slope of 2 points to a purely biphotonic character, doubling pump pulse power causes 4-fold increase in radical signal) and three photons for 540 nm excitation (Figure 4b). The detailed effects of multiphoton excitation on the different excited states (formation quantum yield and lifetime) can be assessed by multi-exponential analysis, as discussed in the next section.

### Global analysis of transient absorption data in the linear and nonlinear excitation regime

For the 540 nm datasets, 4 exponential components convoluted with a Gaussian-shaped pulse of 110 fs (FWHM) and an offset for long lived photoproducts (> 10 ns) are required to obtain a quality fit of all kinetic traces. These components provide 4 characteristic time constants that can be used to describe the behaviour of the different excited states (S_2_, ICT, S_1_, S*) and the long-lived photo-product for P_1_, populated in the end of our time window (1 ns). As already mentioned, for 470 nm excitation, one additional component is needed, which is denoted as S^∼^ (see Figure S8). For all datasets, the first component around 100 fs is associated with S_2_ state, and also includes growth of S_1_, ICT, S*, overlapped with artefacts such as stimulated Raman and eventually any ultrafast Intramolecular Vibrational Relaxation (IVR).

The decay associated spectra (5 exponential components for 470 nm and 4 exponential components for 540 nm excitation) for low energy excitation, 0.4 µJ at 470 nm and 0.8 µJ at 540 nm, are shown in Figure 5a and Figure 5b. Kinetic traces of representative wavelengths with their fits and residuals are given in Figures S9 and S11. The decay associated spectra for both excitations represent S_2_ (yellow), ICT (blue), S_1_ (red), S* (green), S^∼^ (cyan, only for 470 nm excitation) and P_1_ (magenta) species (Figure 5) with associated time constants around 0.6 ± 0.08 ps, 2.5 ± 0.36 ps, 7.0 ± 2.3 ps, 80 ± 30 ps (retrieved time constants are provided in Table 1, standard errors calculated from bootstrapping distributions in Tables S2 and S3). For both excitations, the S* state has a lifetime of about 7 ps with a spectrum characterized by a maximum at 655 nm and a shoulder at 585 nm. The S^∼^ state, which has a longer lifetime of about 80 ps, has its absorption maximum at 570 nm and is similar in shape to the S* state reported by Konold *et al*.^*9*^ (λ_exc_=475 nm). This component is not observed for 540 nm excitation. Global analysis with 4 exponential components (Figures S14 and S15) was also performed for the 470 nm excitation datasets (see the comparison of DAS for 4 and 5 exponential components in Figure S15), however a structure in residuals appears between 10 and 100 ps (Figure S14), and the resulting DAS of the S* state has significantly different properties compared to the ones obtained after 540 nm excitation. In other words, the addition of the S^∼^ component for the 470 nm dataset is indispensable to obtain a consistent description of results recorded with both excitations. Note that time constants retrieved from 4 exponential fit of the 470 nm excited dataset are in agreement with those reported earlier by Konold et al.^9^ suggesting that this component was already present, but not identified, in their experiments carried out on the C-tagged protein using 475 nm excitation at 0.4 µJ excitation energy.

**Table 1:**
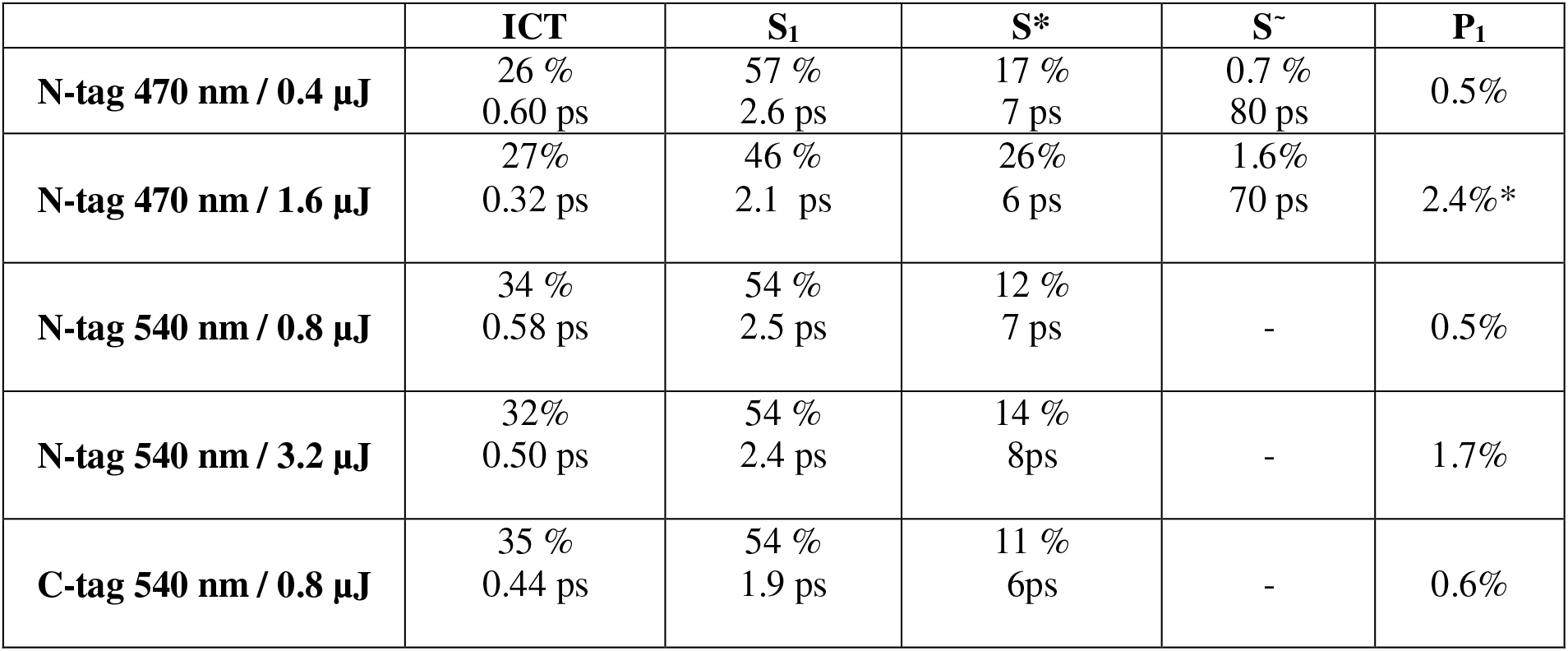
Lifetimes of ICT/S_1_/S*/S^∼^/P_1_ states and estimated formation quantum yields from preexponential factors at 490 nm (See SI for details and Table S1). The following standard errors were estimated (See SI for details, Table S2 and S3).: τ_ICT_ ± 80 fs, τ_S1_ ± 0.36 ps, τ_S*_ ± 2.3 ps, τ_S∼_ ± 30 ps, A_ICT_ ± 5 %, A_S1_ ± 1.9 %, A_S*_ ± 5 %, A_S∼_ ± 1 %, A_P1_ ± 0.1 %. *Note that for P_1_ yield is corrected for its positive absorbance contribution at 490 nm (Figure S4) but not for interfering radical species at high power.

**Figure 5:**
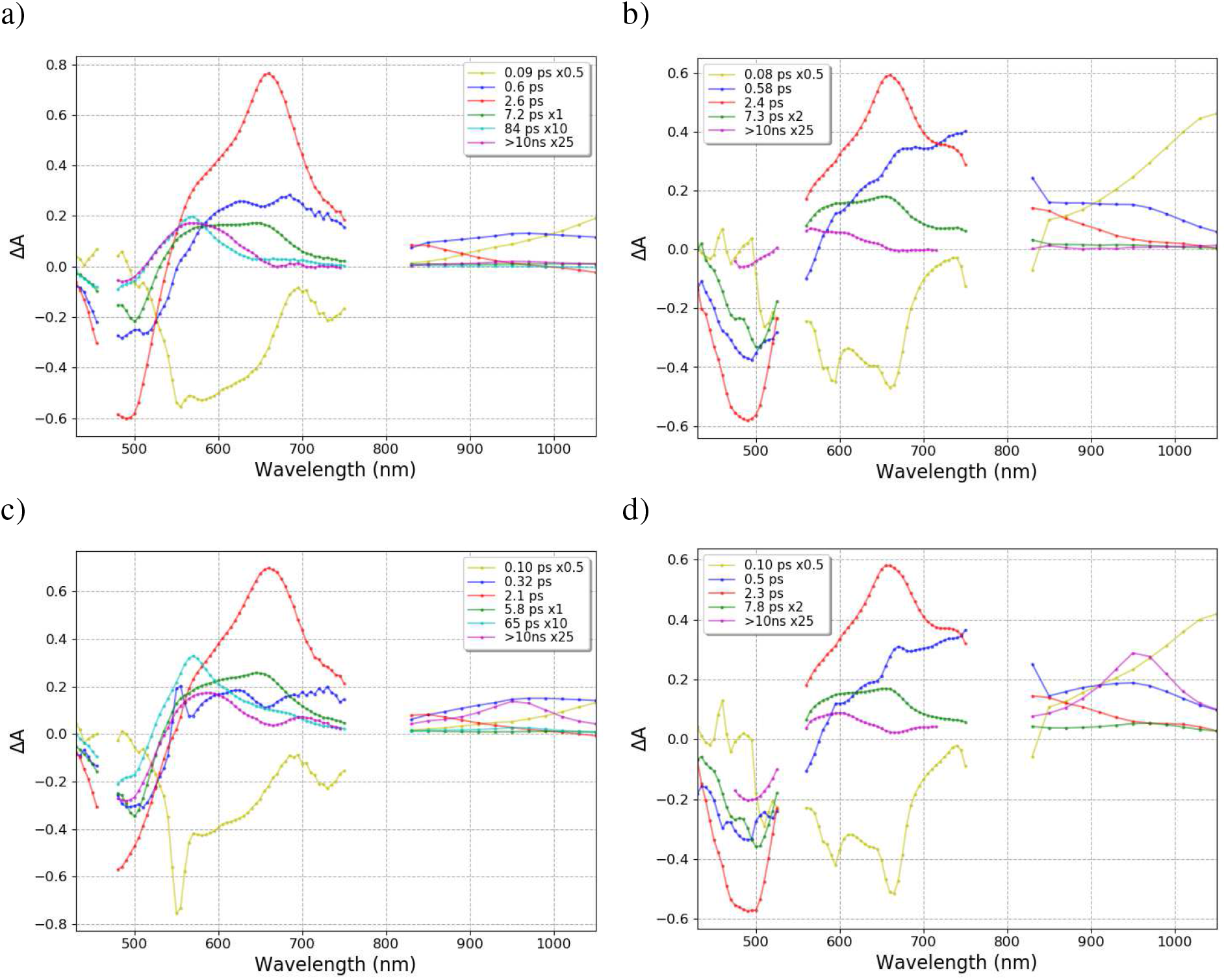
Decay Associated Spectra (DAS) obtained from the global fit of transient absorption data recorded for N-tagged OCP with excitation at a) 470 nm, 0.4 µJ, b) 540 nm, 0.8 µJ, c) 470 nm, 1.6 µJ, d) 540 nm, 3.2 µJ. DAS shown in green are multiplied by 2 for 540 nm excitation and DAS shown in yellow and in magenta are multiplied by 0.5 and 25 for both excitations.

The detailed analysis of the DAS extracted from data collected at low excitation energies (0.4 µJ at 470 nm and 0.8 µJ at 540 nm) reflects the differences in amplitude and shape that were already observed in the raw data. (i) For ICT and S_1_ states the value of absorbance above 700 nm is higher for 540 nm excitation. This results was already reported in the literature,^14-15, 28^ and can be explained by a more pronounced charge transfer character of the S_1_ and ICT states formed after 540 nm excitation (which is clearly visible in Figure 2). (ii) The absorbance of the S* state at 655 nm is about two times higher for λ_exc_=470 nm. (iii) The absorbance at 663 nm for the S_1_ state is substantially higher for λ_exc_=470 nm while for the bleaching extremum (490 nm) it is almost the same for both excitations. (iv) A similar remark can be made for the maximum absorbance of P_1_ (> 10 ns DAS) at 565 nm, higher for 470 nm excitation, while at 490 nm the amplitude is similar for both excitations, pointing to a similar depletion of the OCP^O^ state. However, it needs to be noted that the comparison of the magnitude of the P_1_ positive absorption bands is difficult due to the scattered laser contribution in the 540 nm excitation dataset.

As pointed above P_1_ formation quantum yields can be determined by using the GSB of OCP^O^ depopulation band at 490 nm. Similar method can be used to determine yields of the excited states, which can be estimated from pre-exponential factors at 490 nm (associated with OCP^O^ recovery, Table S1), divided by sum of these excluding P_1_ and S_2_ (see SI for details and Table S1), with results shown in Table 1. It should be underlined here that such approximation assumes that (i) S_1_, ICT and S* are formed from S_2_ in parallel paths (Scheme 2), (ii) these states decay mainly to S0 without any interconversion and (iii) their ESA is small at 490 nm. The comparison of 470 nm excitation (0.4 µJ) and 540 nm excitation (0.8 µJ) indicates a higher and lower contribution of the ICT (34% vs 26%) and S* state (12% vs 17%), upon excitation at 540 nm excitation. Meanwhile, the S_1_ and P_1_ formation quantum yields are independent of the excitation wavelength.

For the high energy excitations datasets, DAS for 1.6 μJ at 470 nm and 3.2 μJ at 540 nm are shown in Figures 5c and 5d, respectively (kinetic traces for representative wavelengths with their fit and residues are given in the Supporting Information, Figures S10 and S12). While the time constants for the S_1_/S*/S^∼^ states do not change with increasing excitation energy, the lifetime of the ICT state slightly decreases with a more pronounced effect observed for 470 nm excitation. The absorbance of positive contribution for excited states does not evolve significantly with increase of the excitation energy. The most relevant feature is that the use of high energy severely increases the negative contribution of the long-lived photo-products (> 10 ns, DAS), which is a mixture of P_1_ and radical cation.

### Influence of tagging on OCP photophysics

The above results point to simpler photo-induced dynamics upon 540 nm excitation (no S^∼^) than 470 nm excitation. Therefore, to study the influence of the His tag, only 540 nm excitation experiments at low energy (0.8 µJ) were performed. Transient spectra for C-tagged OCP-ECN after 540 nm excitation are shown in Figure 6 with DAS (4 exponential components; representative kinetic traces with their fits and residuals are shown in Figure S13). The comparison with N-tagged transient spectra (Figure 2b) shows an increase in the initial absorbance at 740 nm together with a decrease in that at 663 nm (0.2 ps time delay, Figure 6a). Moreover, if one compares the transient spectra at specific time delays after 0.5 ps, or the raw kinetic profiles of C-tagged vs N-tagged OCP (Figure S17), it appears that excited states decay significantly faster in C-tagged OCP. This observation is confirmed by extraction of DAS (see Table 1). Examination of the DAS (Figure 6b) furthermore suggests that C-tagged OCP is characterized by (i) a higher contribution at 740 nm in the S_1_ state, (ii) a different spectral signature of the S* state, with a smaller peak at 655 nm, and (iii) a lower positive signal of P_1_ (Figure S16). Nevertheless, the ratios between intermediate states are quite similar for N-tagged and C-tagged OCP. Despite these differences, the P_1_ formation quantum yield (Table 1) seems to be roughly similar for both N-tagged and C-tagged OCP, and at worst slightly higher for C-tagged OCP. A lower positive band for P_1_ that contributes less at 490 nm (Figure S16), can explain the slight increase in GSB amplitude.

**Figure 6:**
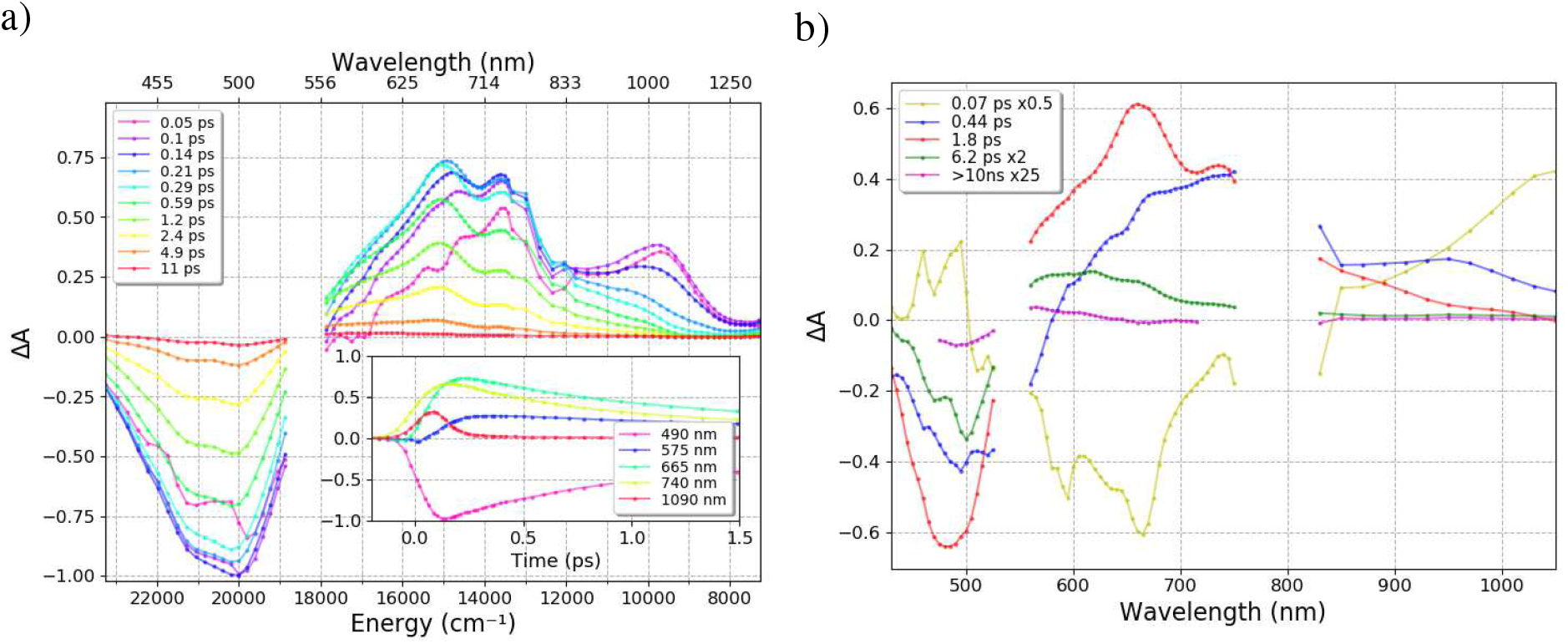
a) Transient absorption spectra between 0.05 and 11 ps for C-tagged OCP excited at 540 nm (0.8 µJ). b) Decay Associated Spectra (DAS) obtained from the global fit of transient absorption data. All datasets were normalized to -1 at bleaching extremum (in both spectral and temporal dimensions, multiply by 0.095 to get original signal). Inset shows evolution of the signal in 1.5 ps time window.

## Discussion

In the present paper, we undertook a detailed photo-physical study on ECN-functionalized *Synechocystis* PCC 6803 OCP with view to investigate the effect on photo-induced excited state dynamics and overall activation yield of (i) excitation at either 470 nm or 540 nm, (ii) His-tagging at the N- (N-tagged) or C- (C-tagged) terminus, and (iii) the excitation power. Our results show that all three factors largely influence OCP excited state dynamics and/or activation yield. Our data also question whether or not the hypothesis that S* is the precursor of P_1_ (and therefore OCP^R^) is correct. Indeed, we offer demonstration that an additional excited state exists upon excitation at 470 nm, and that a radical cation forms upon excitation at high energies – two points that were largely overlooked in the first study which proposed a link between S* and P_1_.^9^ The present work should thus allow a finer understanding of OCP-embedded keto-carotenoid excited state dynamics, and may thereby open avenues towards generation of more efficient OCP.

Most unexpected was the finding that his-tagging at the N- or C-terminus influences the photoactivation speed and excited state dynamics. Proteins are nowadays often expressed recombinantly, with N- or C-terminal extensions that facilitate their purification by affinity chromatography, and OCP is no exception. In nearly all recent studies, a His-tag was accordingly added at the N- or C-terminus, aiding purification and avoiding degradation of the protein.^1, 8^ It should be noted that in the case of OCP, both the N-terminal and C-terminal helices appose on the same face of CTD β-sheet, which serves as the regulatory domain (Scheme 1b). Here, we show that the location of the His-tag affects (i) the photo-activation speed, as derived from initial slopes of OCP^R^ formation (monitored using ΔA at 550 nm) upon continuous irradiation (slower for C-tag, Figure 1b); and (ii) the lifetime of excited states (shorter for C-tag). The population ratio between ps-lived excited states and the P_1_ yield, are yet unaffected by the change in position of the six-histidine tag (Table 1). It is unclear how presence of the latter alters the lifetime and spectral signature of the excited states, however most likely seems the hypothesis that these are affected by different ground state populations of dark-adapted OCP, which each leads to different spectral signatures (notably for S*) and lifetimes for light-induced excited states. Regardless, the most important information is probably that presence of the tag does not influence the P_1_ formation quantum yield (Table 1). Therefore, we attribute the effect of the tag on the photo-activation speed to molecular events occurring on the nano- and millisecond time scales, *i*.*e*. related to carotenoid translocation and/or domain separation (P_1_ to OCP^R^, Scheme 1a).^10-11^ Future studies could address this hypothesis by determining the influence of his-tagging and its location on the outcome of nanosecond flash photolysis experiments.

Since his-tagging slightly modifies the ultrafast dynamics of the OCP, comparison of with earlier studies is only valid if the investigated proteins were tagged at the same location, and ideally with the same tag. Ours results can thus be compared to those obtained by Konold et al.^9^, who studied N-tagged OCP-ECN with 475 nm excitation and were the first to characterize and link the S* and P_1_ states of OCP. To fit their excited states transient spectra, they employed a 3-time-constant multi-exponential model accounting for three ps-lived excited state species, viz. S_1_, ICT, S*. Analogous analysis of the dataset obtained upon 470 nm excitation provides similar results, including a species characterized by a DAS with a maximum at 570 nm and a time constant close to 29 ps (Figure S15). This species was assigned to S* by Konold et al. (24 ps in the article^9^), and likewise by us. However, these results cannot be easily compared with those we obtained upon 540 nm excitation, where S* is characterized by a DAS peaking at 655 nm and a time constant of 7 ps. In contrast, a recent publication from Yaroshevich et al.^13^ reports an S* lifetime of 5.13 ps, in reasonable agreement with the 7 ps lifetime derived from our data collected upon 540 nm excitation. The use of an additional time constant (extra S^∼^ component) to fit data collected upon 470 nm excitation results in an S* signature that is nearly identical to that obtained after 540 nm excitation (maximum of DAS at 655 nm and lifetime of about 6 ps, figure S15). The S^∼^ state is characterized by a DAS peaking at 570 nm and a decay time around 80 ps (Figure 5a). While accounting for only 1% of the excited state signal upon excitation at 470 nm (Table 1, Table S?), addition of S^∼^ to the fitting scheme leads to a higher consistency of determined DAS and time-constants for the 470 nm and 540 nm datasets (Figure S15).

The nature and origin of the extra S^∼^ species is not straightforward to understand. S^∼^ lives longer than usually observed excited states (slightly shorter than 10 ps), and is absent upon 540 nm excitation. A similar state denoted as S^‡^ has been observed in β-carotene, which is characterized by the presence of a 65 ps-lived component located at the high energy edge of ESA. The S^‡^ state was found to not form upon excitation at red edge of β-carotene stationary absorption spectrum, which makes it similar to S^∼^.^38^ The origin of this state was clarified in a later study where it was shown that this component disappears after extensive sample purification.^39^ It was concluded that it must originate from minor impurities of blue-absorbing shorter chain carotenoids. Since our data clearly show that the 80 ps component does not lead to formation of P_1_, we believe that S^∼^ in OCP is likely associated with traces of a non-photoactive carotenoid in OCP. Indeed, it was observed that OCPs produced in *E. coli* not only bind ECN (and/or canthaxanthin) but may also contain an unknown carotenoid of MW 548 (1-6%) and in rare cases also traces of β-carotene.^30^

We also addressed the identity of the precursor of P_1_ and the interplay between the S*, ICT and S_1_ excited states. A comparison of the formation quantum yields of ICT and S* for 470 nm and 540 nm excitations shows an anti-correlation (Table 1), suggesting that paths leading to these states compete. Most recently, S* was assigned to a distorted carotenoid geometry and proposed to precede P_1_.^9^ We found using N-tagged OCP that more S* is produced upon excitation at 470 nm than at 540 nm, despite the P_1_ formation quantum yield being the same value of about 0.5% at the two wavelengths (Table 1). Surely, the error on P_1_ formation quantum yield is large due its low value. Experiments aimed at exploring excitation power dependency showed that 470 nm excitation could involve additional long-lived photo-products (S^∼^ and radical). Hence, it is difficult to determine the quantum yield of P_1_ upon 470 nm excitation with high precision. It can also explain the difference with Konold et al. who reported a value of 1.5%.^9^ Furthermore our value (0.5%) is consistent with the OCP^R^ formation quantum yield estimated by Maksimov et al. to be about 0.2% at 200 ns using nanosecond flash photolysis studies.^12^ Indeed Konold et al. showed that at 200 ns only 40% of the population of P_1_ remained, i.e P_1_ yield should be about 0.5%.^9^ In addition as photo-activation with continuous irradiation at either 470 or 528 nm yields the same initial slopes, we posit that the absorbance at 490 nm provides a correct estimation for the P_1_ formation quantum yield (Table 1). Considering that the S* and ICT states have different yield (Table 1) at both excitation wavelengths, our data thus do not support that S* is the main precursor of P_1_. Rather, they suggest that S_1_ is the most probable P_1_ precursor with a formation quantum yield which is similar for blue and green excitation (Table 1).

## Conclusions

In this work, the effects of excitation energy and wavelength on OCP photoactivation and excited state dynamics were investigated in detail, as well as that of his-tagging at the N- and C-termini. Covering the NIR range (800-1400 nm) in addition to the visible spectrum, we were able to uncover the existence of a carotenoid cation radical, characterized by positive absorbance bands at 960 nm and 700 nm, and whose formation also contributes to the negative depopulation band of OCP^O^. This radical cation is the dominant photoproduct at 1 ns for energies exceeding the linear photoexcitation, calling for a careful consideration of excitation energies in future transient spectroscopy studies on OCP. Excitation at 540 nm compared to 470 nm is better suited to reduce formation of this off-pathway radical species, as a three-photonic regime requires a much higher photon density in the pump pulse. This finding is of importance for the determination of the P_1_ formation quantum yield using the OCP depopulation band. The formation quantum yield of P_1_ is close to 0.5% when the excitation power is low enough to avoid formation of the cation radical (0.3 mJ/cm^2^ or less for λ_exc_=470 nm, 0.6 mJ/cm^2^ or less for λ_exc_=540 nm, Figures S5 and S6), consistent with previous estimates from nanosecond flash photolysis studies.^12^ Another important result regarding 470 nm excitation is the existence of a previously unnoticed S^∼^ species (≈80 ps lifetime), which does not form upon excitation at 540 nm and display similar spectral features as P_1_. Most likely, this S^∼^ species originates from impurities, possibly β-carotene. Accounting for this additional species observed upon 470 nm excitation, global fitting of the datasets for the two excitation wavelengths gives similar DAS and lifetime for ICT/S_1_/S* regardless of the excitation wavelength (470 nm vs 540 nm). Our analysis shows that the S* lifetime is about 7 ps for 470 nm and 540 nm excitations. Because the S^∼^ component most likely originates from a chromophore not involved in the photo-activation of OCP, modelling the dynamics using data obtained with green excitation is more relevant for the photo-dynamics of OCP, particularly since photo-activation experiments using continuous irradiation do not show dependence on the excitation wavelength. An important result is that the formation quantum yield for S* and ICT differs for both excitations (Table 1), while the P_1_ and S_1_ formation quantum yield do not change. This strongly supports the hypothesis that S_1_, but not S*, is the main precursor of P_1_. We thus suggest a new path for the picosecond photo-dynamics leading to P_1_ - photoactivation of OCP (from Synechocystis N-tagged with echinenone), using results upon 540 nm excitation and assuming parallel formation and decay of picosecond states (ICT, S_1_, S*) with yields estimated from OCPO depopulation band (Scheme 2, Table 1)

**Scheme 2:**
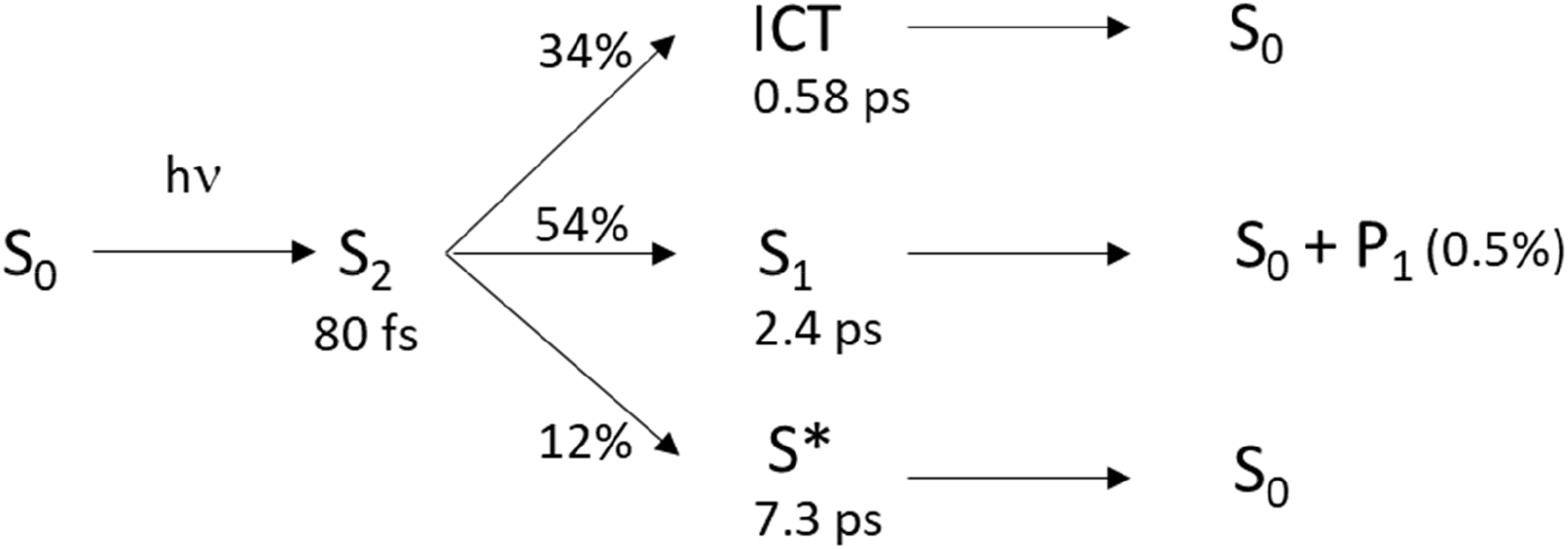
Picosecond photo-dynamics of Synechocystis N-tagged echinenone-OCP upon 540 nm excitation (yields are estimated from GSB recovery of OCPO at 490 nm, Table 1)

Finally, the comparison of ps dynamics in C-tagged vs N-tagged OCP revealed significant differences in the DAS and formation quantum yields of S*, and an overall faster relaxation of excited states in C-tagged OCP. Also, the photo-activation speed observed under continuous irradiation was found to differ significantly. Yet, the P_1_ yield was unchanged. Hence we pose that tagging of OCP rather influences molecular events occurring past 1ns, i.e. carotenoid translocation and/or structural changes (Scheme 1a). Further studies conducted on the nanosecond-millisecond range could verify this hypothesis.

## ACKNOWLEDGMENTS

This work was performed with financial support from the Polish National Science Centre (NCN), project 2018/31/N/ST4/03983, and the French National Research Agency (grant ANR-18-CE11-0005 to M.S., D.K., I.S. and J.P.C.).

## SUPPORTING INFORMATION

Supplementary materials includes details on “normal” and “redshifted” sub-populations existing in the dark-adapted state, pump pulse excitation, time resolved spectra at high excitation energy, analysis of kinetics traces, comparison of P_1_ spectra upon different excitation vs His-tag and evaluation of error using bootstrapping method.

