## Supplementary materials for "A unifying perspective of the ultrafast photo-dynamics of Orange Carotenoid Protein from *Synechocystis*: peril of high-power excitation, existence of different S* states and influence of tagging"

### Contents

1) Steady state UV-vis absorption spectra: page 2-3

2) Transient absorption spectra of N-tagged OCP in the visible range: page 4

3) Determination of P1 formation quantum yield: page 4-5

4) Pump pulse characterization: page 6-7

5) Transient absorption spectra at high excitation energy: page 7

6) Analysis of kinetics traces: page 8-12

7) $P_1$ signature: page 13

8) Estimation of formation quantum yields from pre-exponential factors at 490 nm: page 13-14

9) Estimation of the error on time constant and pre-exponential factor: 14-15

10) Comparison of kinetic profile of C-tagged and N-tagged OCP: page 16

### 1) Steady state UV-vis absorption spectra

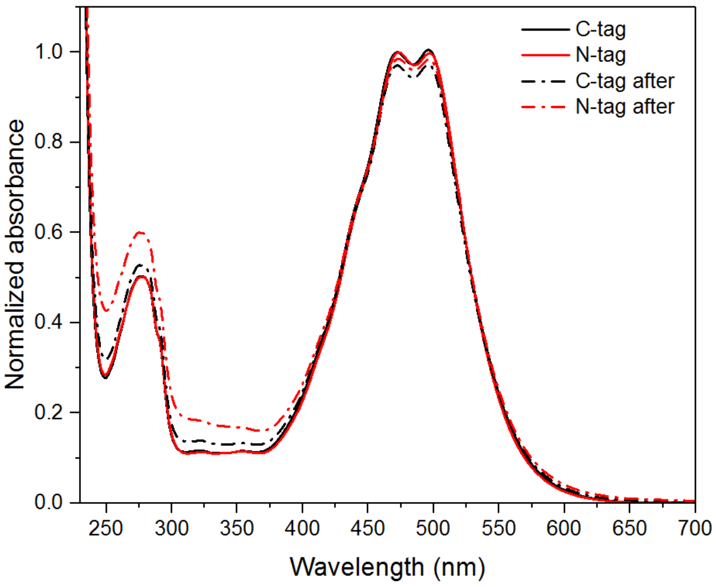


**Fig. S1 Absorption spectra of dark-adapted state (OCP$^{o}$) for N-tagged and C-tagged OCP, with spectra recorded after one of the femtosecond transient absorption experiments.**

**a)**

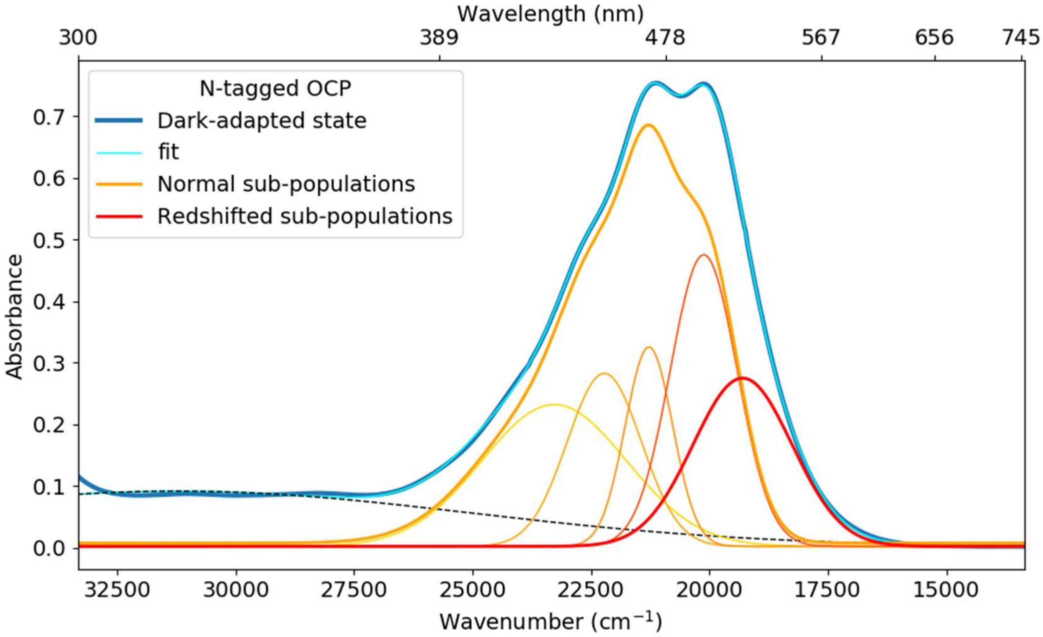


**b)**

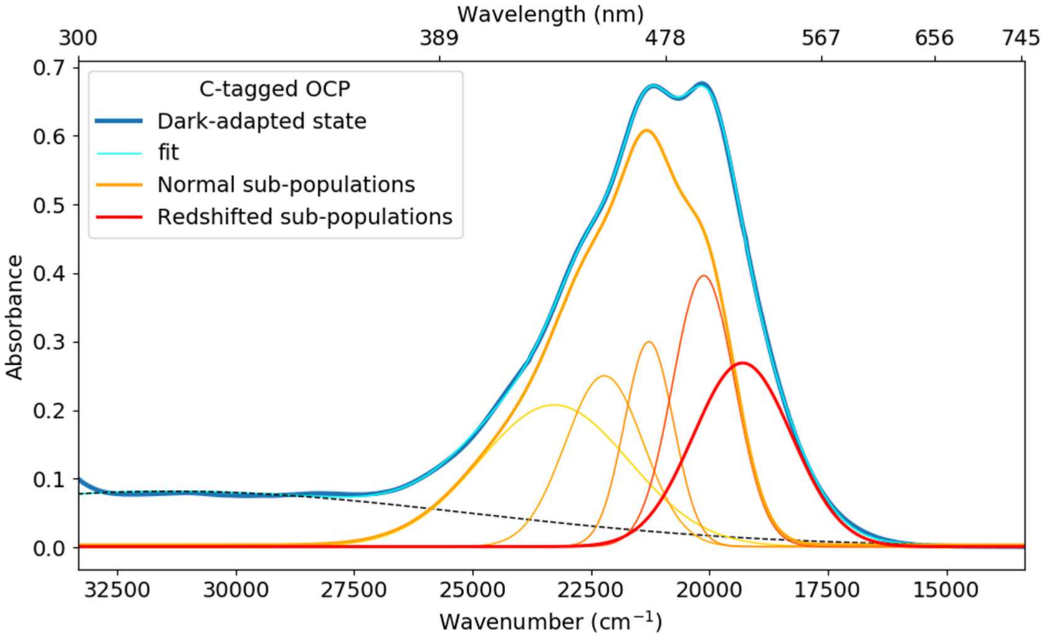


**Fig. S2 Gaussian-sum decomposition of the absorption spectra of a) C-tagged OCP b) N-tagged OCP into "normal" and "redshifted" sub-populations existing in equilibrium in the dark-adapted state (OCP$^{O}$).**

### 2) Transient absorption spectra of N-tagged OCP in the visible range

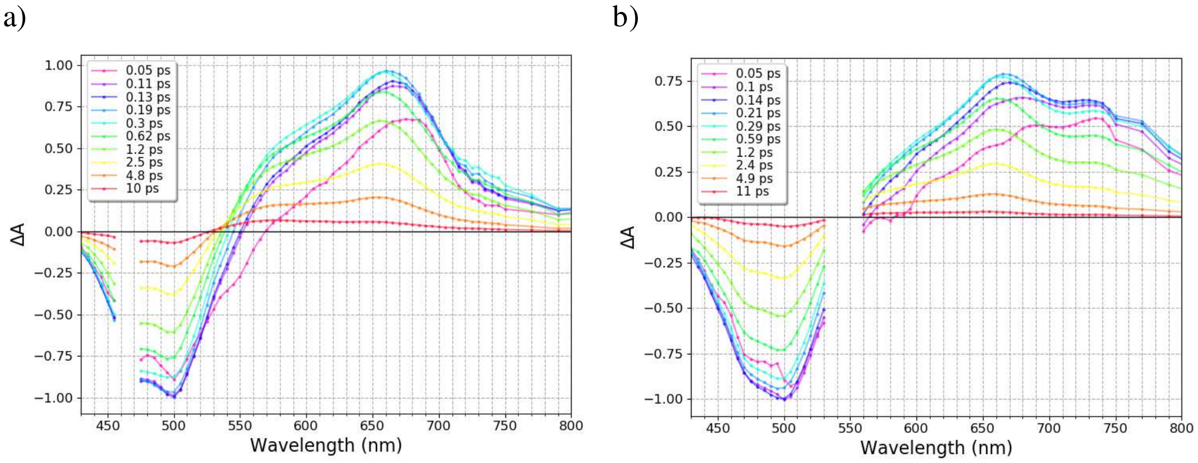


**Fig. S3 Visible region of the transient absorption spectra between 0.05 and 10 ps for N-tagged OCP-ECN excited at a) 470 nm (0.4 µJ), and b) 540 nm (0.8 µJ). All datasets were normalized to -1 at the bleaching extremum (in both spectral and temporal dimensions).**

### 3) Determination of P1 formation quantum yield

Formation quantum yield of $P_1$ was estimated from the comparison of initial bleached OCP population and final $P_1$ ΔA(490nm) signal extracted from DAS decomposition. To estimate concentration of the initially excited population, transient spectrum at 0.1 ps time delay was used and -63 000 $M^{-1}cm^{-1}$ extinction coefficient was assumed at the extremum of the depopulation band at 490 nm [1] (normalized to -1). In order to calculate concentration of the $P_1$ state, amplitude of the DAS for $P_1$ at 490 nm (Figure 5) was used with an extinction coefficient at 490nm of -29 500$M^{-1}cm^{-1}$. This coefficient was estimated by firstly calculating $OCP^R$ minus $OCP^O$ differential spectrum (Figure 1a), assuming an extinction coefficient of 63 000 $M^{-1}cm^{-1}$ for $OCP^O$ at 490 nm (Figure S4). Konold et al. used these spectra to compensate for induced absorbance of $P_1$ at 490 nm [2]. However, despite having similar spectral signatures, $OCP^R$ difference absorption has a maximum at 550nm, while $P_1$ one is peaking around 565 nm (Figure S4). Therefore we shifted stationary $OCP^R$ spectrum to the red by 30nm and then calculated difference spectrum, in a such a way to get a maximum at 565nm for positive signal and a zero crossing point similar to the one of $P_1$ DAS. Difference spectrum of the $OCP^R$ state (measured data, red curve) and calculated one for $P_1$ (redshifted before subtraction, black curve) is shown in the Figure S4, in comparison with $P_1$ DAS obtained with both used excitations.

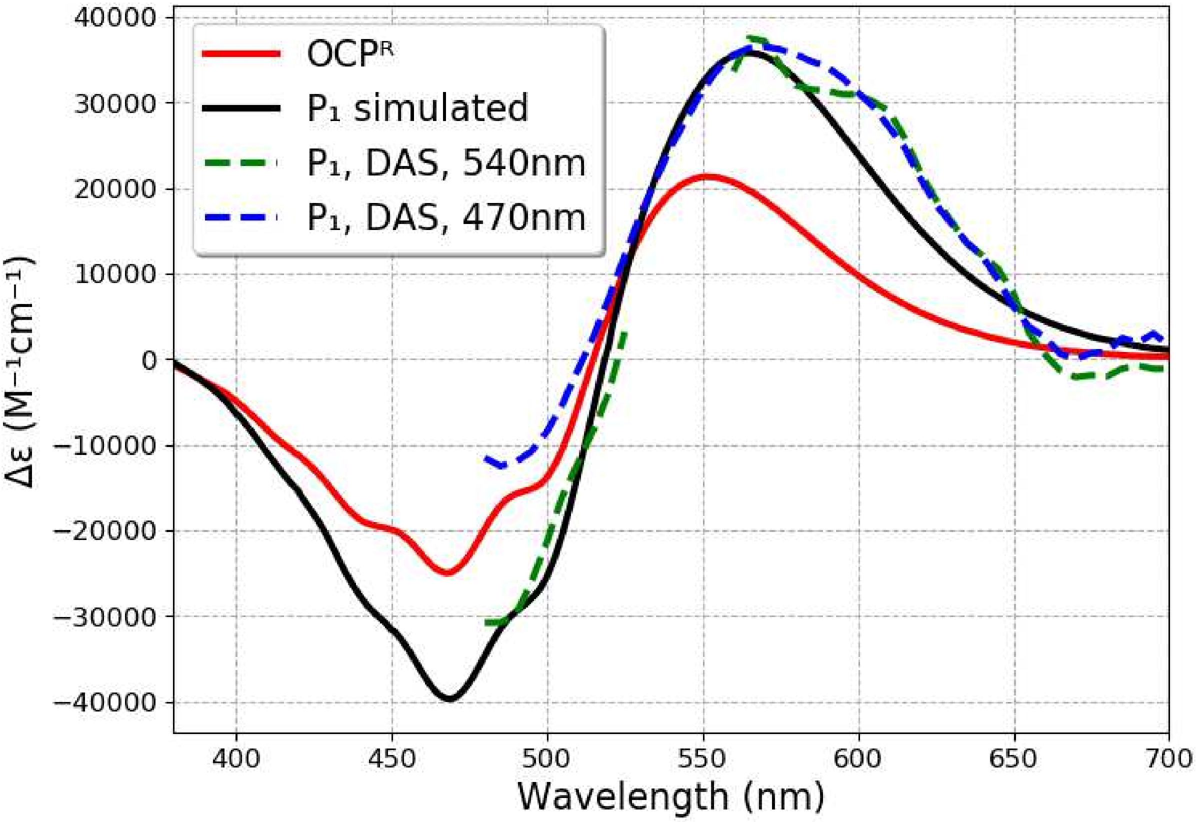


**Fig.S4 : Difference spectrum of $OCP^R$ state minus $OCP^O$ (measured data, red curve, Figure 1a), difference spectrum calculated one for $P_1$ by redshifting $OCP^R$ stationary spectrum (black curve), DAS for $P_1$ (Figure 5) upon 540 nm excitation (0.8 μJ, green dashed line) and upon 470 nm excitation (0.4 μJ, blue dashed line)**

### 4) Pump pulse characterization

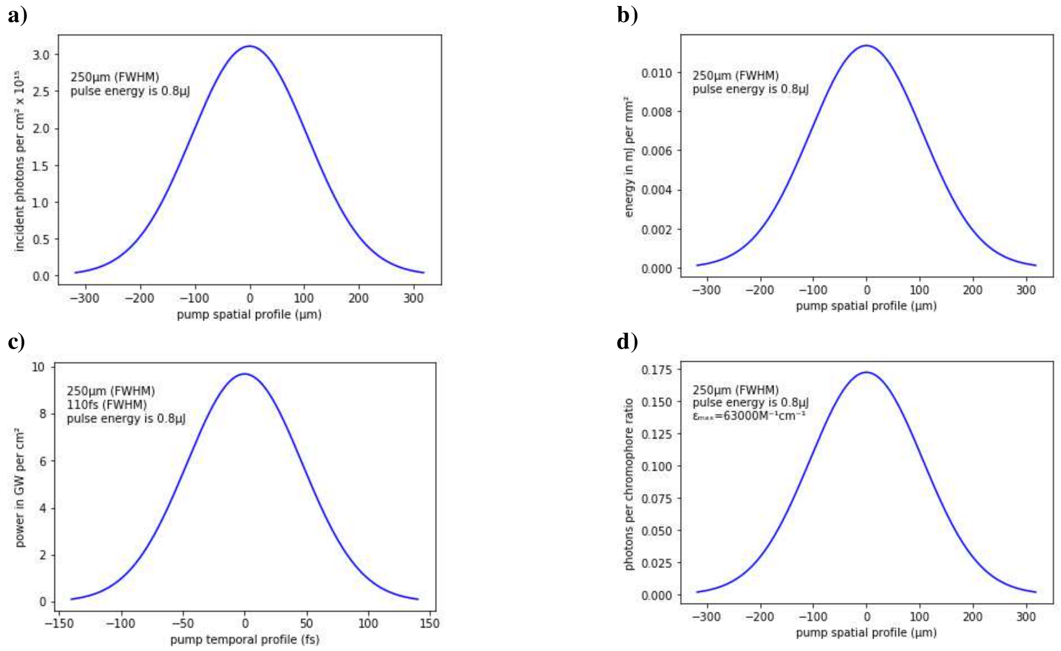


**Fig. S5 Simulations showing the energy distribution (a) photons per cm$^2$, b) mJ.mm$^{-2}$, c) GW.cm$^{-2}$, d) photons absorbed per chromophore) of the pump pulse for N-tagged OCP experiments, excitation 540 nm, energy 0.8 µJ, sample absorption A(540 nm, 2 mm thickness) = 0.19**

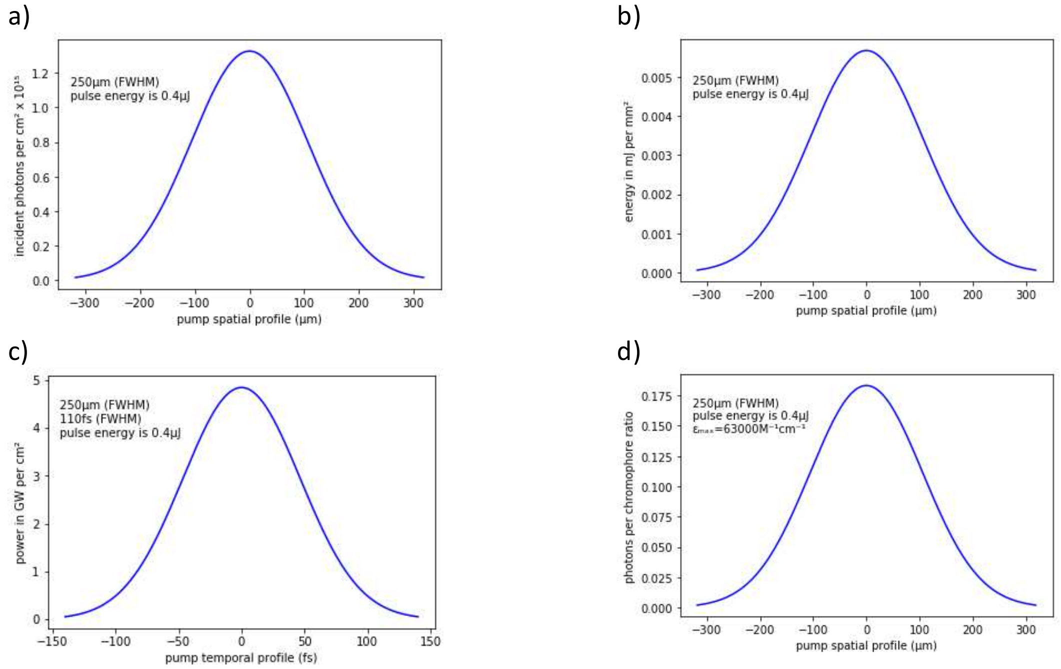


**Fig. S6 Simulations showing the energy distribution (a) photon per cm$^2$, b) mJ.mm$^{-2}$, c) GW.cm$^{-2}$, d) photon absorbed per chromophore) of the pump pulse for N-tagged OCP experiments, excitation 470nm, energy 0.4uJ, sample absorption A(470 nm, 2 mm thickness) = 0.49**

Method to determine number of photons absorbed per pulse and ratio of echinenone excited in the excited volume: the number of photons in the pump pulse per unit surface of the plane perpendicular to the pump beam was calculated from the measured pump energy divided by the energy of a photon at the excitation energy. Next a 2D Gaussian distribution defined on the cuvette (cylinder of a 2D Gaussian base and 2 mm length) was assumed at this point, with measured FWHM (250 µm), the same on both axes. The derived distribution was multiplied by $(1 - 10^{-A_{exc}})$ to obtain the number of absorbed photons per unit surface in the excited volume. The OCP concentration was calculated using a molar absorption coefficient of $\epsilon$ = 63 000 $M^{-1}cm^{-1}$ at the absorption maximum accordingly to Sluchanko et al. [1] Then number of OCP molecules excited in the illuminated volume was calculated as concentration multiplied by the Avogadro constant and the cuvette internal thickness. Finally the ratio of OCP excited per pulse in the pump volume was found from number of absorbed photons divided by the number of OCP molecules.

### 5) Transient absorption spectra at high excitation energy

a) b)

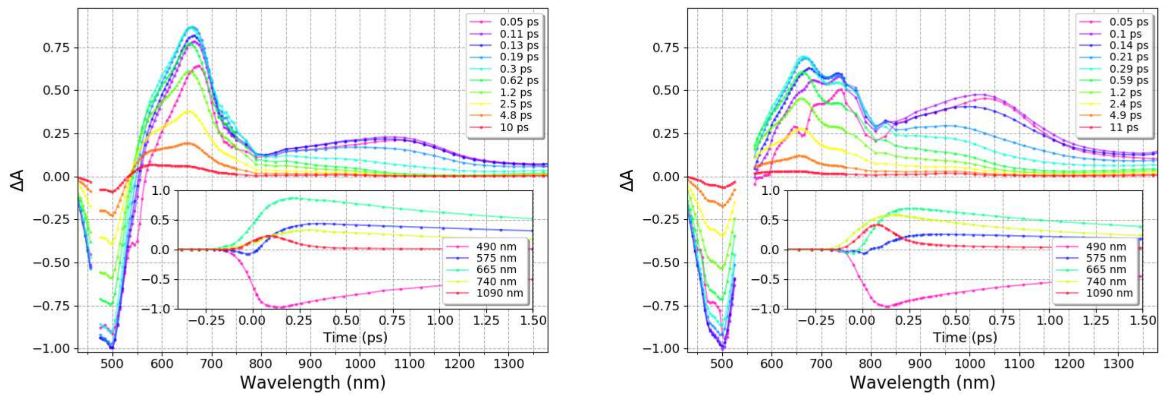


**Fig. S7 Transient absorption spectra between 0.05 and 10 ps of N-tagged OCP-ECN excited at a) 470 nm (1.6 µJ, 110 fs FWHM, 250 µm FWHM), and b) 540 nm (3.2 µJ, 110 fs, 250 µm FWHM). All datasets were normalized to -1 at bleaching minimum (in both spectral and temporal dimensions). Insets show the growth of the signal.**

### 6) Analysis of kinetic trace analysis

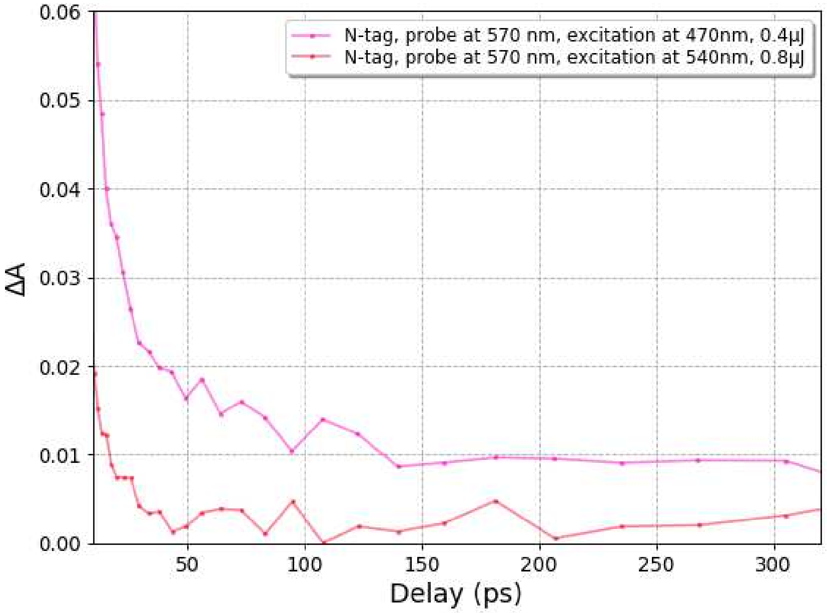


**Fig. S8: Kinetics at 570 nm for excitation at 470 nm and 540 nm, respectively.**

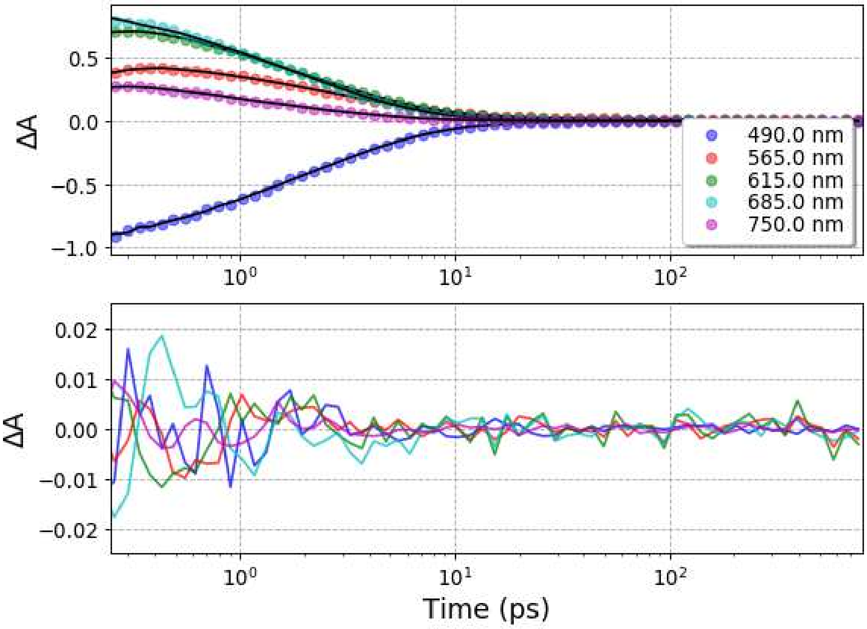


**Fig. S9 Representative kinetic traces with their fit and residuals of global fit analysis done on N-tagged OCP (5 exponential components convoluted with a Gaussian-shaped pulse of 110 fs (FWHM) and an offset for long lived photoproducts (> 10 ns)), excitation at 470 nm, 0.4 µJ**

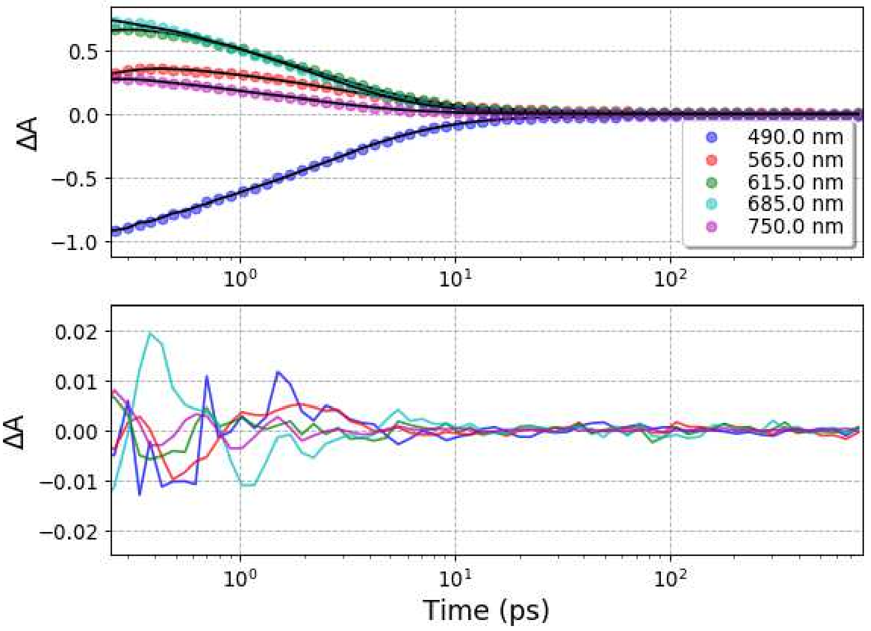


**Fig. S10 Representative kinetic traces with their fit and residuals of global fit analysis done on N-tagged OCP (5 exponential components convoluted with a Gaussian-shaped pulse of 110 fs (FWHM) and an offset for long lived photoproducts (> 10 ns)), excitation at 470 nm, 1.6μJ**

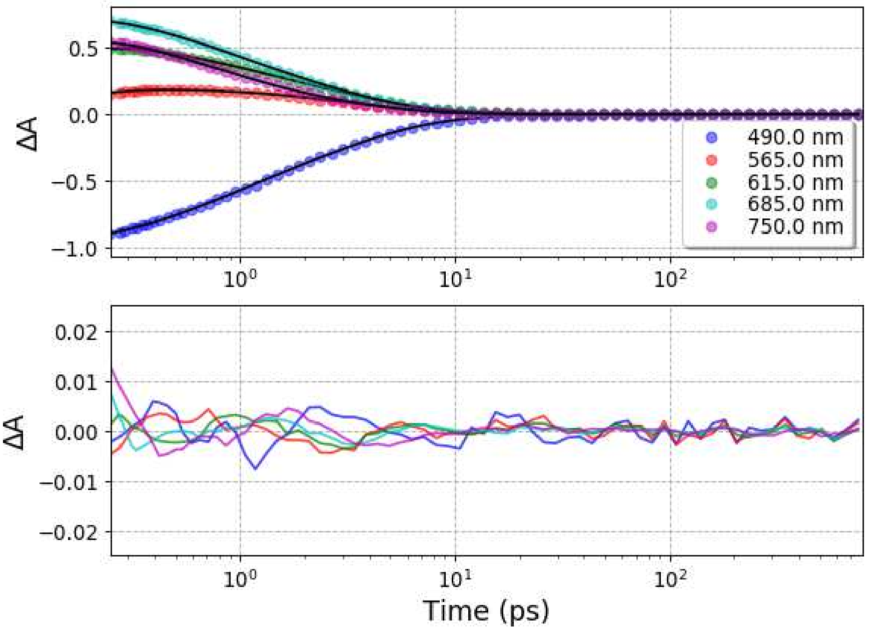


**Fig. S11 Representative kinetic traces with their fit and residuals of global fit analysis done on N-tagged OCP (4 exponential components convoluted with a Gaussian-shaped pulse of 110 fs (FWHM) and an offset for long lived photoproducts (> 10 ns)), excitation at 540 nm, 0.8 μJ**

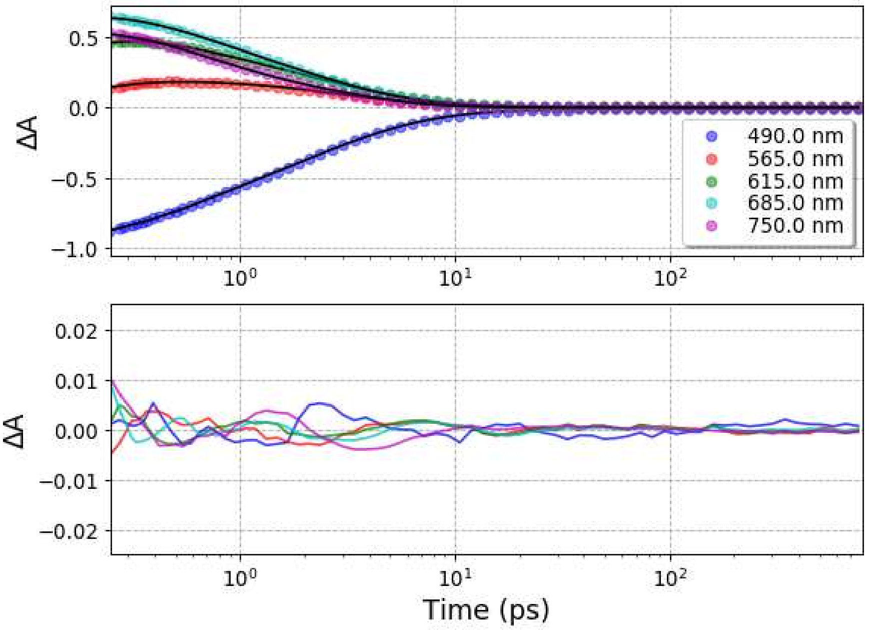


**Fig. S12 Representative kinetic traces with their fit and residuals of global fit analysis done on N-tagged OCP (4 exponential components convoluted with a Gaussian-shaped pulse of 110 fs (FWHM) and an offset for long lived photoproducts (> 10 ns)), excitation at 540nm, 3.2µJ**

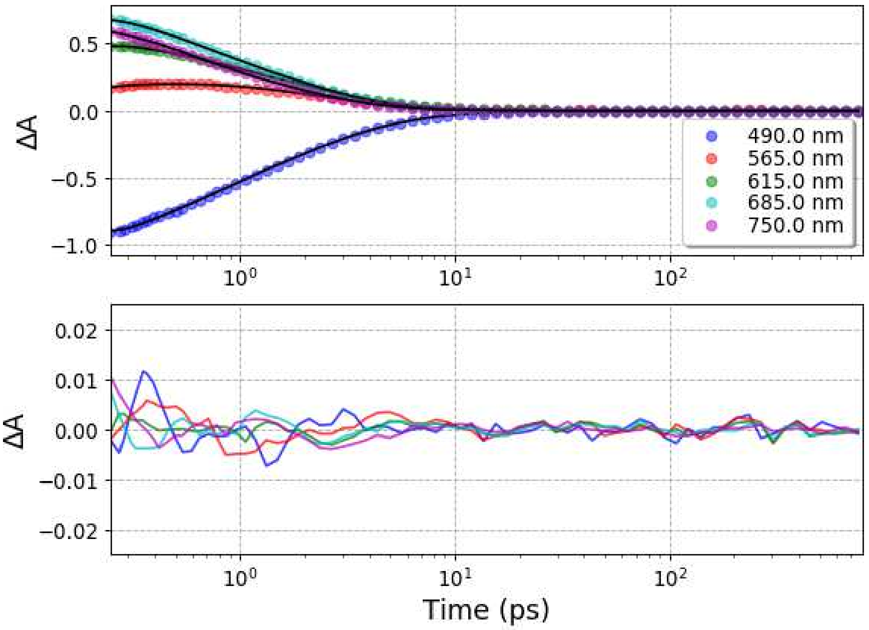


**Fig. S13 Representative kinetic traces with their fit and residuals of global fit analysis done on C-tagged OCP (4 exponential components convoluted with a Gaussian-shaped pulse of 110 fs (FWHM) and an offset for long lived photoproducts (> 10 ns)), excitation at 540 nm, 0.8 µJ**

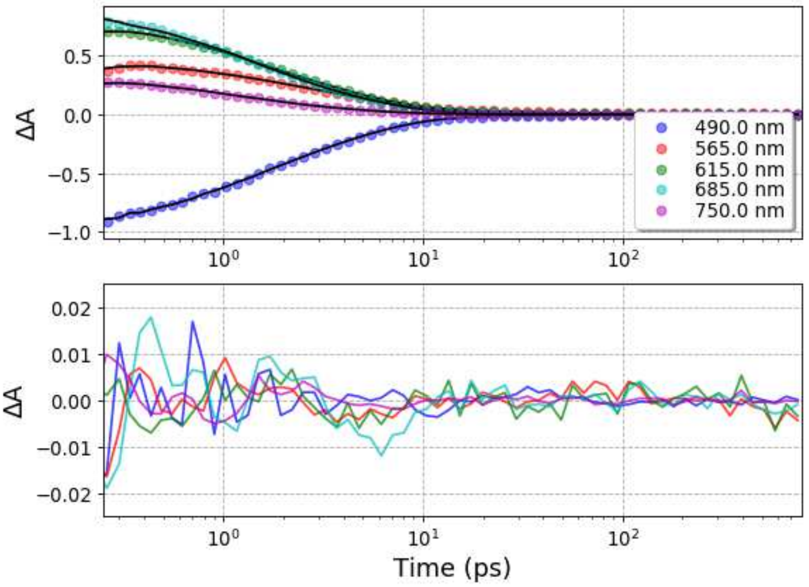


**Fig. S14 Representative kinetic traces with their fit and residuals of global fit analysis done on N-tagged OCP (4 exponential components convoluted with a Gaussian-shaped pulse of 110 fs (FWHM) and an offset for long lived photoproducts (> 10 ns)), excitation at 470 nm, 0.4 µJ**

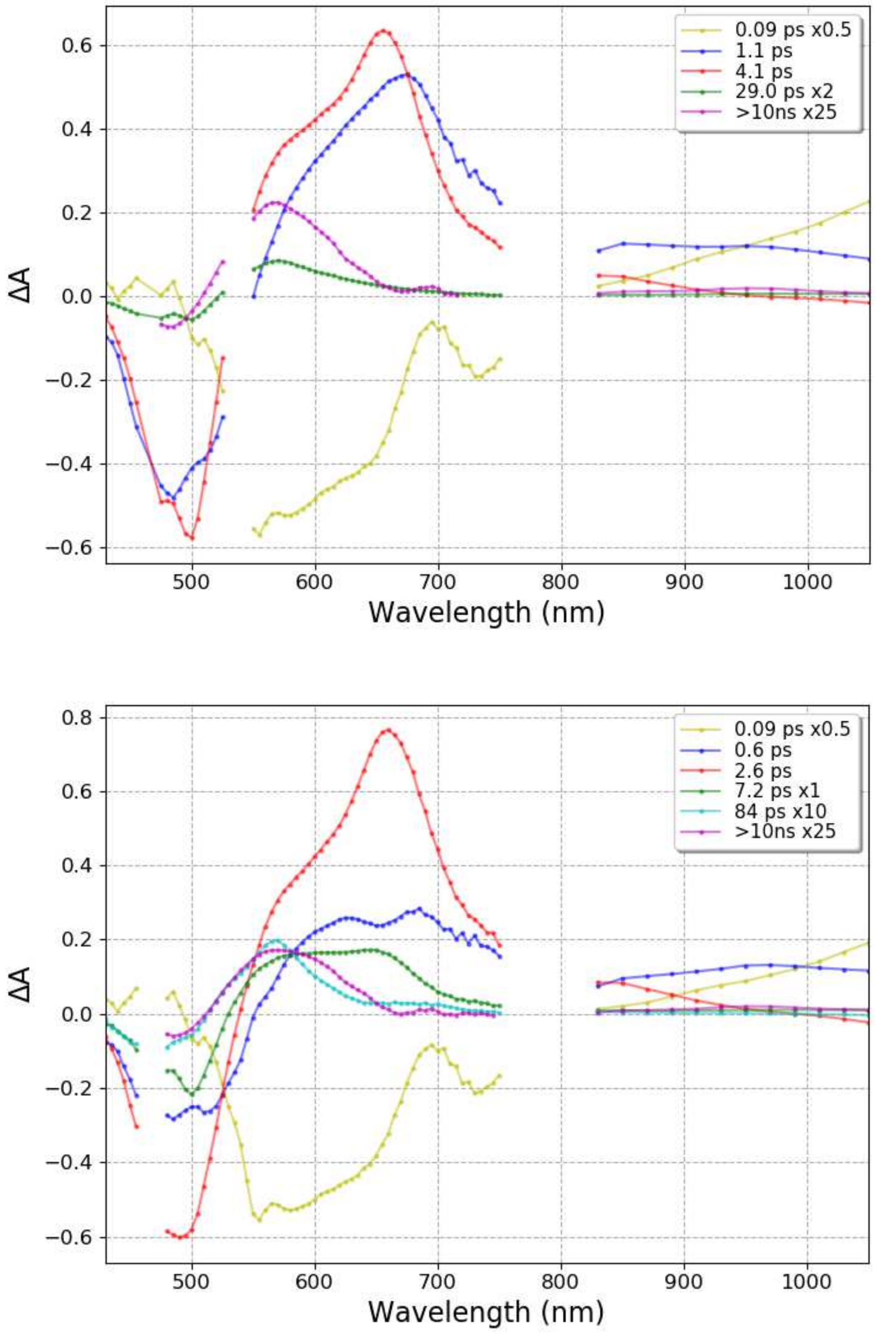


**Fig. S15 Decay Associated Spectra (DAS) obtained from the global fit (top panel 4 exponential components convoluted with a Gaussian-shaped pulse of 110 fs (FWHM) and an offset for long lived photoproducts (> 10 ns), bottom panel 5 exponential components convoluted with a Gaussian-shaped pulse of 110 fs (FWHM) and an offset for long lived photoproducts (> 10 ns)) of transient absorption data recorded of N-tagged OCP with excitation at 470 nm (0.4 µJ).**

### 7) $P_1$ signature

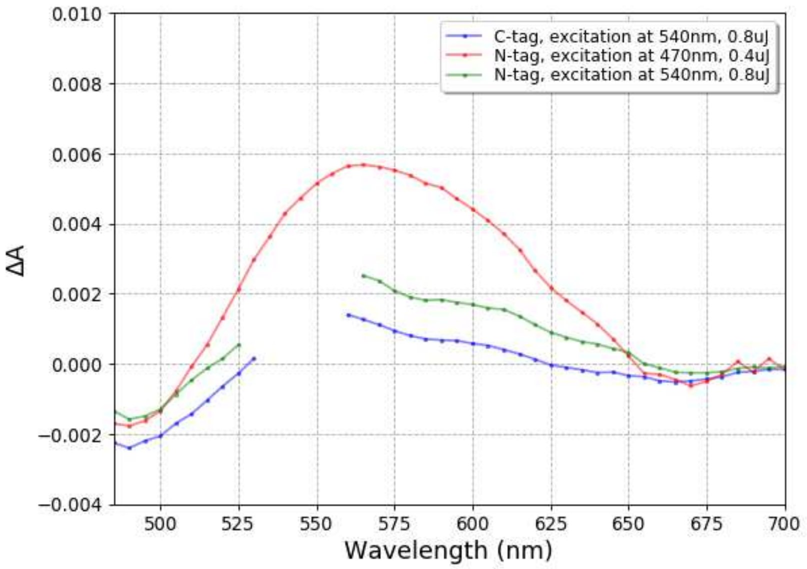


**Fig. S16 Transient absorption spectra averaged around 1 ns ($P_1$ state) of N-tagged and C-tagged OCP.**

### 8) Estimation of formation quantum yields from pre-exponential factors at probe 490 nm

The formation quantum yield of $S_1$, ICT or S* are estimated from pre-exponential factors at probe 490 nm, i.e. their DAS value at 490 nm (Figure 5, Figure 6, Table S1), divided by sum of $S_1$, ICT and S* amplitudes excluding $P_1$ and $S_2$ (see equations below), with results shown in Table 1 and Scheme 2. It should be underlined here that such approximation assumes that (i) $S_1$, ICT and S* are formed from $S_2$ in parallel paths, (ii) these states decay mainly to $S_0$ without any interconversion and (iii) their excited state absorption is small at 490 nm. For the $P_1$ formation quantum yield its positive absorbance contribution at 490 nm is estimated and taken into account (Figure S4).

Table S1: DAS value at 490 nm for ICT, S1, S*, S~ and P1.

| | ICT | S1 | S* | S~ | P1 |
|---|---|---|---|---|---|
| N-tag 470 nm / 0.4 µJ | 0.27 | 0.60 | 0.17 | 0.0070 | 0.0023 |
| N-tag 470 nm / 1.6 µJ | 0.30 | 0.53 | 0.29 | 0.018 | 0.011 |
| N-tag 540 nm / 0.8 µJ | 0.37 | 0.58 | 0.13 | - | 0.0023 |
| N-tag 540 nm / 3.2 µJ | 0.33 | 0.57 | 0.15 | - | 0.0082 |
| C-tag 540 nm / 0.8 µJ | 0.41 | 0.63 | 0.13 | - | 0.0028 |

Equations to calculate formation quantum yield for ICT/$S_1$/S*/$P_1$:

$$\Phi_{ICT} = DAS_{ICT}(490nm)/\left(DAS_{ICT}(490nm) + DAS_{S_1}(490nm) + DAS_{S*}(490nm)\right)$$

$\Phi_{S_1} = DAS_{S_1}(490nm)/\left(DAS_{ICT}(490nm) + DAS_{S_1}(490nm) + DAS_{S*}(490nm)\right)$

$\Phi_{S*} = DAS_{S*}(490nm)/\left(DAS_{ICT}(490nm) + DAS_{S_1}(490nm) + DAS_{S*}(490nm)\right)$

$\Phi_{P_1} = DAS_{P_1}(490nm) \cdot 63000/29500$

where $DAS_X(490nm)$is DAS value of X state at 490nm, extracted from data normalized to -1 at the bleaching extremum (Figure 5 and Figure 6).

### 9) Estimation of the error on time constants and pre-exponential factors

To assess the error of the extracted time constants and quantum yields (Table 1 in the main text), bootstrap analysis was employed. For the sake of simplicity, we used one dataset. We selected N-tagged OCP, done with excitation at 540 nm, 0.8 μJ, as the most representative one for this study.

Our bootstrap procedure consists of the following steps:

a) Dataset was fitted using a procedure described in Materials and Methods.
b) Residuals from the obtained fit were used to calculate the standard deviation (estimation of the noise in the data) for each kinetics separately (noise levels are different depending on spectral region). Delays below 0.3 ps were excluded due to strong contribution of artifact signals such as SRA (where residuals don't reflect noise, but they have shape of the artifacts).
c) Sampled datasets were generated, where additional random noise was simulated from standard deviations obtained in previous step, using gaussian distribution, and added to the original data. 6473 sample datasets were generated in total.
d) Each generated sample dataset was fitted using the same procedure as for the original fit. It allowed to build a distribution of each fit parameter, and calculate a standard errors.

Described procedure yields errors and distributions shown in tables S2 and S3 (below). Obtained sets of errors are expected to be similar between datasets. These results do not contain information about error of the S˜ state, because 540 nm excited dataset do not contain S˜ state. Nevertheless, it is clear that errors for quantum yield and lifetime of the S˜ state are huge, due to very small contribution. Our fitting procedure on 470 nm excited dataset usually returned something between 50 ps and 110 ps for the S˜, depending on initial fit conditions and tuning of the fitting procedure itself. So we assume that error of the S~ is better than +/- 30 ps.

**Table S2: Distributions of parameters obtained from bootstrap procedure of N-tagged OCP (excitation 540 nm, 0.8 µJ, 6473 samples). Note that amplitudes are calculated as preexponential factor divided by sum of preexponential factors excluding $S_2$ state.**

| | ICT | S1 | S* | P1 |
|---|---|---|---|---|
| τ /ps | 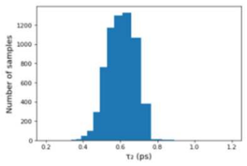 | 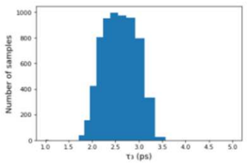 | 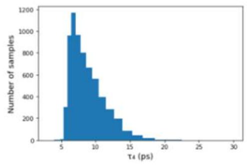 | |
| $A_{490\ nm}$ | 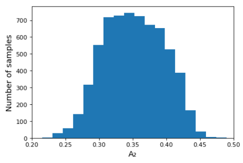 | 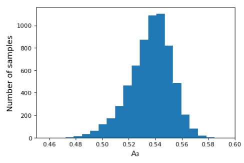 | 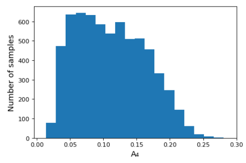 | 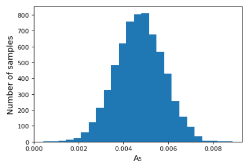 |

**Table S3: Standard errors calculated from bootstrapping distributions of N-tag (exc. 540 nm, 0.8 µJ, Table S1)**

| | ICT | $S_1$ | S* | $P_1$ |
|---|---|---|---|---|
| σ(τ) / ps | 0.076 | 0.36 | 2.3 | - |
| σ($A_{490\ nm}$) | 0.046 | 0.019 | 0.052 | 0.0011 |

### 10) Comparison of kinetic profiles of C-tagged and N-tagged OCP

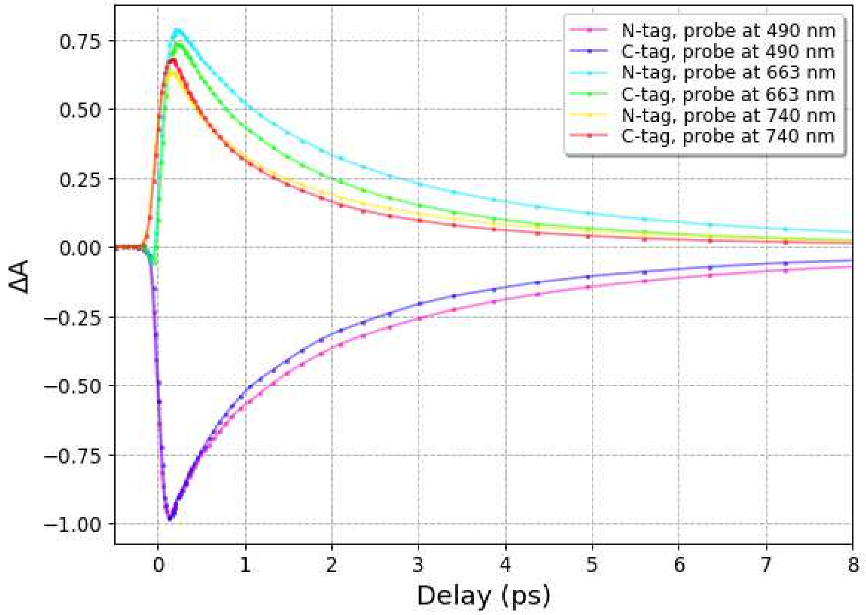


**Fig. S17 Comparison of kinetic profiles of C-tag and for N-tag (540 nm, 0.8 μJ)**
